# Enhanced MicroRNA Accumulation and Gene Silencing Efficiency through Optimized Precursor Base Pairing

**DOI:** 10.1101/2025.09.04.674230

**Authors:** Juan-José Llorens-Gámez, Pedro José García-Cano, Sara Rico-Rodrigo, Lucía Duyos-Casanova, Sara Toledano-Franco, Alberto Carbonell

**Author notes:** Corresponding author: Alberto Carbonell.

## Abstract

MicroRNAs (miRNAs) are endogenous 21-nucleotide small RNAs that direct sequence-specific silencing of complementary messenger RNAs to regulate a wide range of biological processes. In plants, miRNA precursors are processed from imperfect foldback structures by the RNase III enzyme DICER-LIKE1, in coordination with accessory proteins. While mismatches flanking the miRNA/miRNA* duplex in endogenous precursors can strongly influence miRNA accumulation, their impact has not been thoroughly examined in the context of artificial miRNAs (amiRNAs) used for targeted gene silencing in plants. Here, using silencing sensor systems in *Nicotiana benthamiana*, we systematically investigated how base-pairing at or near DCL1 cleavage sites affects amiRNA production from the recently described minimal *shc* precursor. Independent pairing of naturally mismatched positions revealed that introducing a G-C pair immediately upstream of the mature amiRNA remarkably enhances amiRNA accumulation and silencing efficiency. This effect was further validated in Arabidopsis transgenic lines targeting endogenous genes and confirmed by deep sequencing, which revealed highly accurate processing and predominant release of the intended amiRNAs, supporting the specificity of the approach. Our findings show that a single structural modification in an amiRNA precursor can significantly enhance the efficacy of amiRNA-mediated gene silencing. This optimized amiRNA platform is well suited for large-scale functional genomics screens and should facilitate the development of next-generation crops with enhanced resilience to environmental stresses.

**SIGNIFICANCE STATEMENT:** A large-scale mutational analysis of how base-pairing at or near DCL1 cleavage sites affects artificial microRNA (amiRNA) production from the minimal *shc* precursor revealed that introducing a single G-C base pair immediately upstream of the first DCL1 cleavage site significantly boosts amiRNA accumulation and gene-silencing efficiency. The incorporation of this structural tweak into a high-efficiency amiRNA platform provides an optimized, highly specific RNAi tool for functional genomics and crop engineering.

## INTRODUCTION

MicroRNAs (miRNAs) are ∼21-nucleotide (nt) non-coding small RNAs (sRNAs) that guide ARGONAUTE (AGO) proteins to complementary messenger RNAs (mRNAs), leading to target cleavage or translational inhibition. In plants, miRNAs regulate genes encoding transcription factors and other proteins involved in critical biological processes including development, stress responses and hormone signaling (Bologna and Voinnet, 2014; Zhan and Meyers, 2023). They originate from longer precursors transcribed by RNA polymerase II, termed primary miRNAs (pri-miRNAs), which fold into characteristic stem-loop foldback structures recognized and sequentially cleaved by the DICER-LIKE 1 (DCL1) endoribonuclease in the nucleus [reviewed recently in (Yu *et al*., 2025)]. A miRNA duplex of approximately 21 nt that features 2-nt 3′ overhangs is excised, and typically one strand—the guide miRNA—is incorporated into an ARGONAUTE (AGO) protein, where it functions to direct gene silencing by base pairing with complementary RNA targets (Fang and Qi, 2016; Carbonell, 2017b).

Plant miRNA precursors are highly diverse in size but share a conserved structural architecture, typically comprising a ∼15–17 bp basal stem (BS), a central miRNA/miRNA* duplex, and a distal stem-loop (DSL) region that varies in length and conformation and is bordered by single-stranded regions (Bologna *et al*., 2009; Cuperus *et al*., 2011). In the predominant base-to-loop processing pathway, the DICER-LIKE1 (DCL1) enzyme–alongside cofactors SERRATE (SE) and HYPONASTIC LEAVES1 (HYL1)–first cleaves the precursor at the basal stem to generate a shorter hairpin intermediate. A second cleavage, positioned approximately 21 nt from the initial site, releases the miRNA/miRNA* duplex, which possesses characteristic 2-nt 3′ overhangs and is subsequently stabilized by 2′-O-methylation via the methyltransferase HUA ENHANCER1 (HEN1) (Zhu *et al*., 2013; Song *et al*., 2010; Werner *et al*., 2010; Mateos *et al*., 2010). Alternatively, some miRNA precursors follow an alternative loop-to-base processing mode, in which DCL1 initiates cleavage at the terminal loop and proceeds toward the base to excise the miRNA duplex (Bologna *et al*., 2013; Chorostecki *et al*., 2017, p.20178; Addo-Quaye *et al*., 2009; Bologna *et al*., 2009). Despite key structural features for accurate and efficient miRNA processing have been identified (Bologna *et al*., 2013; Moro *et al*., 2018; Zhu *et al*., 2013; Cuperus, Montgomery, *et al*., 2010; Song *et al*., 2010; Werner *et al*., 2010; Mateos *et al*., 2010; Chorostecki *et al*., 2017), the importance of specific sequences in this matter has been largely unknown. Recently, a genome-wide examination of base-pairing interactions at the DCL1 cleavage sites in natural *Arabidopsis thaliana* (Arabidopsis) and eudicot miRNA precursors revealed an enrichment of base pairs among the four positions flanking the miRNA/miRNA* duplex, with sequence biases at specific positions (Rojas *et al*., 2020). Interestingly, the base pairing of naturally occurring mismatches generally altered miRNA accumulation, with the nucleotide identity and position affecting processing efficiency.

Artificial miRNAs (amiRNAs) exploit the native plant miRNA biogenesis pathway to induce targeted gene silencing with high specificity, becoming a versatile tool for plant functional genomics and biotechnology (Cisneros *et al*., 2021). AmiRNAs are engineered 21-nt sRNAs designed *in silico* to reprogram the endogenous miRNA processing and silencing pathways for specific repression of selected target transcripts with minimal off-target effects (Carbonell, 2017a; Ossowski *et al*., 2008; Tiwari *et al*., 2014). AmiRNAs are typically generated *in planta* by expressing endogenous *MIRNA* precursors in which the native miRNA/miRNA* duplex is replaced with the synthetic amiRNA/amiRNA* duplex, thereby producing a functional pri-miRNA precursor processed by the endogenous machinery. Choosing an optimal pri-miRNA backbone is essential to ensure precise and efficient processing of the engineered precursor, and accumulate high amiRNA levels required for effective silencing. The 521-nt long Arabidopsis *MIR390a* (*AtMIR390a*) precursor is processed accurately and efficiently relative to other plant pri-miRNAs commonly used for amiRNA production (Lunardon *et al*., 2021; Carbonell *et al*., 2014), and has been broadly applied for amiRNA expression across various plant species–including both model systems and crops–to achieve effective silencing of endogenous genes and viral RNAs (Lunardon *et al*., 2021; Carbonell *et al*., 2019; Vasav *et al*., 2022; Berbati *et al*., 2023; Kadam and Barvkar, 2024). Recently, the minimal structural and sequence requirements for producing effective amiRNAs from the *AtMIR390a* precursor were systematically analyzed (Cisneros *et al*., 2023). As a result, highly effective and accurately processed amiRNAs were produced form a shorted chimeric “*shc*” precursor of only 89 nt, including the complete BS of *AtMIR390a* (without additional ssRNA segments) and the DSL region derived from *Oryza sativa MIR390* with a 2-nt deletion. Importantly, the *shc* precursor has a compact DSL region of only 15 nt, allowing the synthesis of the entire foldback with just two oligonucleotides. This simple structure has facilitated the development of a cost-effective, high-throughput cloning methodology for direct insertion of amiRNA sequences into a suite of ‘B/c’ vectors incorporating the *AtMIR390a* BS, thereby simplifying and accelerating the production of amiRNA constructs (Cisneros *et al*., 2023).

Here, we used the recently described silencing sensor systems in *Nicotiana benthamiana* (Cisneros *et al*., 2023; Cisneros and Carbonell, 2025) to systematically investigate how base-pairing mismatches at or near DCL1 cleavage sites within the minimal *shc* precursor affect amiRNA biogenesis and function. We functionally screened a large collection of constructs expressing amiRNAs targeting two *N. benthamiana* genes, from modified *shc*-based precursors with distinct base pairing configurations. By combining phenotypic, biochemical and molecular assays, we show that *shc* precursors in which adenine at position 18 is substituted with a guanine (A18G) yield increased amiRNA levels and markedly enhanced silencing. Finally, the superior performance of A18G-modified *shc* precursors was further validated in Arabidopsis transgenic plants expressing amiRNAs against endogenous genes whose silencing induced a visible and quantifiable phenotype. Furthermore, high-throughput sequencing-based analysis of precursor processing, showed that A18G-modified *shc* precursors are accurately processed and release authentic amiRNAs.

## RESULTS

### Enhanced accumulation of miR390a from a modified *AtMIR390a* precursor without mismatches at DCL1 first cleavage site

To assess the impact of eliminating mismatches at the DCL1 initial cleavage site on miR390a biogenesis, we engineered a modified Arabidopsis *MIR390a* precursor in which the adenine at position 18 was substituted with guanine (A18G), thus restoring base pairing with cytosine at position 89 (C89) (Figure 1a). This modified construct (*35S:AtMIR390a-A18G*) and a wild-type control (*35S:AtMIR390a*) were transiently expressed in *N. benthamiana* leaves through Agrobacterium infiltration. Each construct was agroinfiltrated into two leaves per plant across three biological replicates. A *35S:GUS* construct expressing *Escherichia coli* β-glucuronidase *uidA* gene served as a negative control. sRNA blot analysis at 2 days post-agroinfiltration (dpa) revealed a significant increase in miR390a accumulation from the mismatch-corrected precursor compared to the wild-type (Figure 1b).

**Figure 1.**
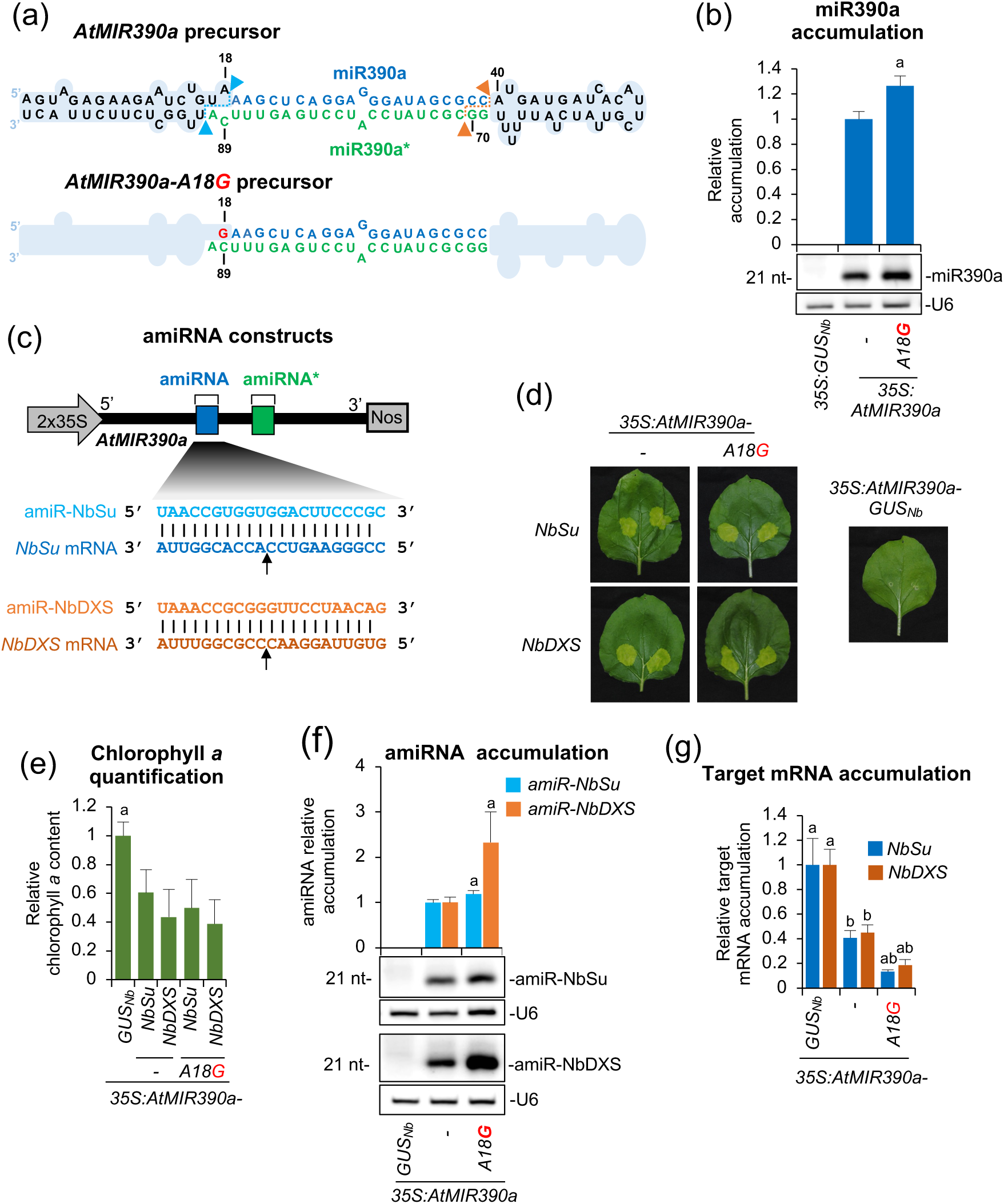
Functional analysis of endogenous and modified Arabidopsis *MIR390a* (AtMIR390a)-based precursors without mismatches at DCL1 first cleavage site. (a) *AtMIR390a* and *AtMIR390A-A18G* foldback diagrams. DCL1 first and second cleavage sites are marked with blue and orange arrows, respectively. *AtMIR390a*, miR390a and miR390a*nucleotides are highlighted in black, blue and green, respectively. Mutated G at position 18 is shown in red. Shapes corresponding to *AtMIR390a* basal stem and distal stem loop are in light blue. (b) Northern blot detection of miR390a in RNA preparations from agroinfiltrated leaves at 2 days post-agroinfiltration (2 dpa). The graph at top shows the mean (*n* = 3) + standard deviation miR390a relative accumulation (*35S:AtMIR390a* = 1). Bar with the letter ‘a’ is significantly different from that of the *35S:AtMIR390a* control sample (P < 0.05 in pairwise Student’s *t*-test comparison). (c) Diagram of *AtMIR390a*-based amiRNA constructs including the base-pairing of amiRNAs and target mRNAs. Nucleotides corresponding to the guide strand of the amiRNA against *NbSu* and *NbDXS* are in blue and orange, respectively, while nucleotides of target mRNAs are in dark blue and orange, respectively. The arrows indicate the amiRNA-predicted cleavage site. (d) Photos at 7 dpa of leaves agroinfiltrated with different constructs. (e) Bar graph showing the relative content of chlorophyll *a* in patches agroinfiltrated with different constructs (*35S:AtMIR390a-GUS* = 1.0). Bars with the letter ‘a’ are significantly different from that of the *35S:AtMIR390a-GUS* control sample (P < 0.05 in pairwise Student’s t-test comparisons). (f) Northern blot detection of amiR-NbSu and amiR-NbDXS in RNA preparations from agroinfiltrated leaves at 2 dpa. Other details are as in (b). (g) Target mRNA accumulation in agroinfiltrated leaves. Mean relative level (n = 3) + standard error of *NbSu* or *NbDXS* mRNAs after normalization to *PROTEIN PHOSPHATASE 2A* (*PP2A*), as determined by quantitative RT-PCR (qPCR) (*35S:AtMIR390a-GUS_Nb_*= 1.0 in all comparisons). Bars with the letter ‘b’ or ‘a’ are significantly different from that of the corresponding control samples *35S:AtMIR390a-GUS_Nb_*or *35S:AtMIR390a-NbSu/35S:AtMIR390a-NbDXS*, respectively (P < 0.05 in pairwise Student’s t-test comparisons).

### Increased amiRNA accumulation and activity from the modified *AtMIR390a-A18G* precursor

To determine whether the *AtMIR390a-A18G* precursor could enhance accumulation of artificial miRNAs (amiRNAs), we used two previously described gene-silencing reporters targeting the endogenous *N. benthamiana SULPHUR* (*NbSu*) and *DXS* (*NbDXS*) genes, which encode magnesium chelatase subunit CHLI and 1-deoxy-D-xylulose-5-phosphate synthase, respectively (Cisneros *et al*., 2023). Visible bleaching phenotypes indicate efficient silencing of these targets. Constructs expressing amiR-NbSu and amiR-NbDXS from the modified precursor were generated (Figure 1c) and agroinfiltrated into two regions per leaf of two leaves across three plants. Parallel infiltrations with constructs expressing amiR-NbSu, amiR-NbDXS and amiR-GUS (an amiRNA targeting *E. coli* β *uidA* gene) (Cisneros *et al*., 2022) from the wild-type *AtMIR390a* precursor were also performed as controls. At 7 dpa, leaf sectors expressing amiR-NbSu or amiR-NbDXS displayed strong bleaching phenotypes (Figure 1d), correlating with significant reductions in chlorophyll *a* content compared to the control (Figure 1e). Next, two leaves of three different plants were independently agroinfiltrated in the whole leaf surface with each of the amiRNA constructs described above. RNA blot analysis at 2 dpa of leaves fully infiltrated with each amiRNA construct confirmed that amiR-NbSu and amiR-NbDXS accumulated at higher levels when expressed from the A18G-modified precursor (Figure 1f). Consistently, RT-qPCR analysis revealed significantly reduced *NbSu* and *NbDXS* transcript levels in samples expressing the corresponding amiRNAs from the modified precursor (Figure 1g), confirming increased silencing efficiency.

### Analysis of amiRNA accumulation in modified *sch* precursors without mismatches at DCL1 first cleavage site

To further investigate whether the base pairing at DCL1 first cleavage site enhances amiRNA accumulation from other precursors, we analyzed the recently described *shc* amiRNA precursor (Figure 2a) (Cisneros *et al*., 2023). Mutations were introduced at positions 18 and 73 of the basal stem to assess the effect on amiRNA accumulation of nucleotide identity and specific base-pair combinations at DCL1 first cleavage site. Constructs expressing amiR-NbSu and amiR-NbDXS from *shc-*based variant precursors including all possible nucleotide combinations at DCL1 first cleavage site (positions 18/73) were generated (Figure 2b). These constructs, along with the GUS-targeting control, were transiently expressed in *N. benthamiana* leaves through agroinfiltration, as previously described.

**Figure 2.**
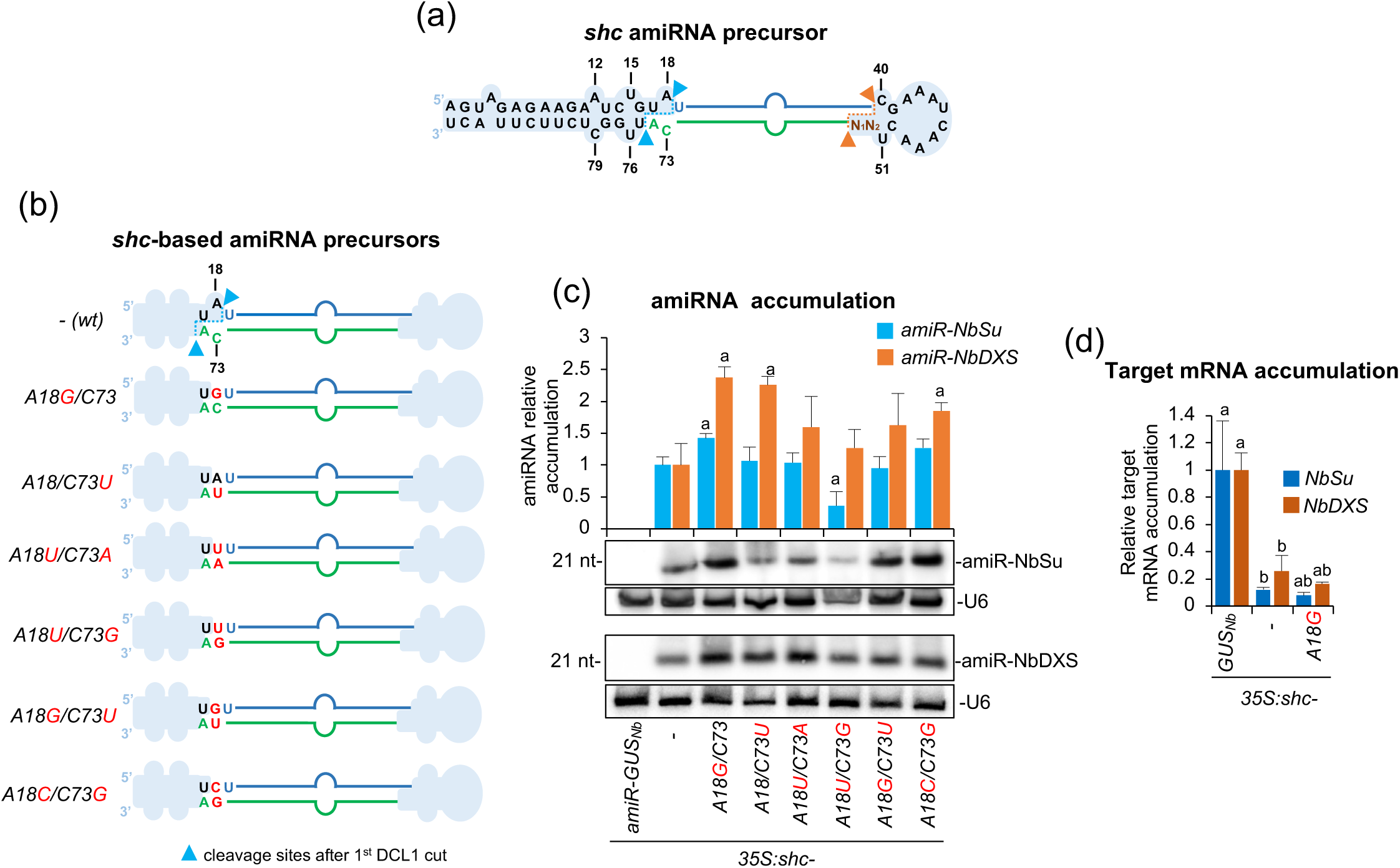
Functional analysis of wild-type and modified *shc-*based precursors without mismatches at DCL1 first cleavage site. (a) *shc* foldback diagrams with DCL1 first and second cleavage sites are marked with blue and orange arrows, respectively. miR390a and miR390a*nucleotides are highlighted in blue and green, respectively. Relevant unpaired positions are numbered. (b) Diagrams of the amiRNA precursors with mutated nucleotides at position 18 in red. Nucleotides of the precursor, amiRNA and amiRNA* are in black, blue and green, respectively. The shapes corresponding to *shc* basal stem or distal stem loop are in light blue. (c) Northern blot detection of amiR-NbSu and amiR-NbDXS in RNA preparations from agroinfiltrated leaves at 2 dpa. Bar with the letter ‘a’ is significantly different from that of the corresponding wild-type *shc-NbSu/shc-NbDXS* control samples (P < 0.05 in all pairwise Student’s *t*-test comparisons). (d) Target mRNA accumulation in agroinfiltrated leaves. Mean relative level (n = 3) + standard error of *NbSu* or *NbDXS* mRNAs after normalization to *PROTEIN PHOSPHATASE 2A* (*PP2A*), as determined by quantitative RT-PCR (qPCR) (*35S:shc-GUS_Nb_*= 1.0 in all comparisons). Bars with the letter ‘b’ or ‘a’ are significantly different from that of the corresponding control samples *shc-amiR-GUS_Nb_*or wild-type *shc-NbSu/shc-NbDXS* samples, respectively (P < 0.05 in all pairwise Student’s *t*-test comparisons).

sRNA blot analyses of RNA samples from agroinfiltrated leaves collected at 2 dpa revealed that all precursor variants produced detectable levels of mature amiRNAs (Figure 2c). Regarding amiR-NbSu, accumulation was significantly higher in samples expressing the A18G/C73 or A18U/C73G variants relative to the wild-type configuration (Figure 2c). In the case of amiR-NbDXS, all modified precursors exhibited enhanced accumulation, although only those with A18G, C73U or A18G/C73G modifications showed statistically significant increases (Figure 2c). Finally, target transcript levels were analyzed in tissues expressing amiR-NbSu or amiR-NbDXS from the A18G/C73 precursor, which produced the highest (and significant) accumulation of both amiRNAs. RT-qPCR analysis revealed a significant reduction in *NbSu* and *NbDXS* transcript abundance in these samples, therefore confirming the enhanced silencing efficiency conferred by this modified precursor (Figure 2d).

### Analysis of amiRNA accumulation in modified *sch* precursors without mismatches at DCL1 second cleavage site or at other basal stem positions

Next, we sought to determine how nucleotide identity and base pairing affect amiRNA accumulation at the DCL1 second cleavage site. For that purpose, we further modified the *shc* precursor at positions 40 and 51, which define the second DCL1 cleavage site (Figure 3a). A series of *shc-*based precursors containing all plausible nucleotide combinations at positions 40/51 were generated (Figure 3a), each expressing amiR-NbSu or amiR-NbDXS. These constructs, along with the GUS-targeting control, were transiently expressed in *N. benthamiana* leaves through agroinfiltration, as previously described. sRNA blot analyses of RNA samples from agroinfiltrated leaves collected at 2 dpa revealed detectable levels of amiR-NbSu and amiR-NbDXS from all modified precursor variants (Figure 3b), with no significant differences among the different variants, indicating that base paring at DCL1 second cleavage site in *shc* has not a significant effect on amiRNA accumulation.

**Figure 3.**
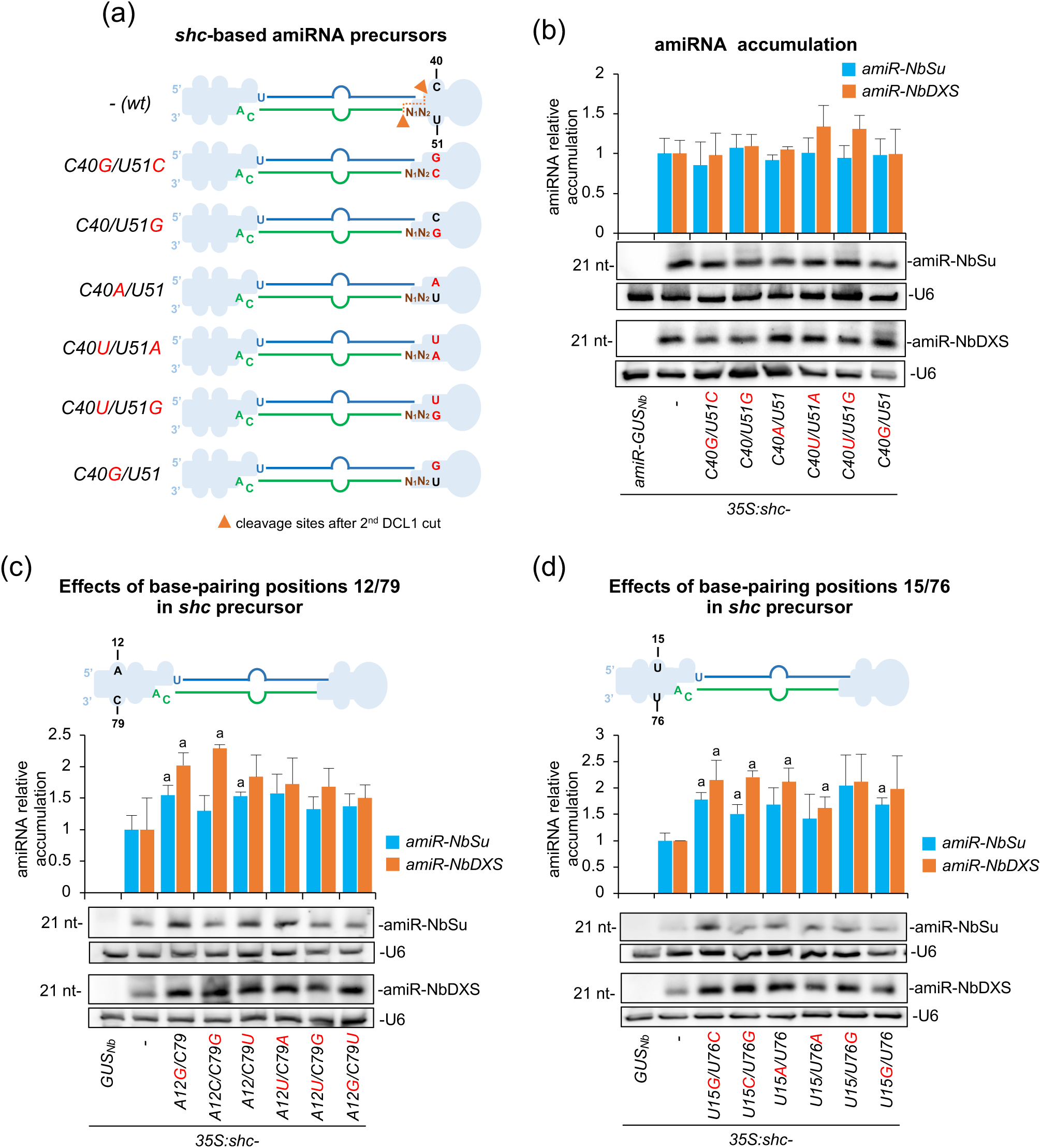
Functional analysis of wild-type and modified *shc-*based precursors without mismatches at DCL1 second cleavage site and at positions 12/79 and 15/76. (a) Diagrams of the amiRNA precursors with mutated nucleotides at position 40/51 (corresponding to DCL1 second cleavage site) in red. Other details are as in Figure 2b. (b) Northern blot detection of amiRNAs. Other details are as in Figure 2c. (c) Top, diagram of wild-type *shc* amiRNA precursor with unpaired position 12/79 highlighted. Other details are as in Figure 2b. Bottom, Northern blot detection of amiRNAs. Other details are as in Figure 2c (d) Top, diagram of wild-type *shc* amiRNA precursor with unpaired position 15/76 highlighted. Other details are as in Figure 2b. Bottom, Northern blot detection of amiRNAs. Other details are as in Figure 2c.

To further dissect how internal stem base pairing influences amiRNA processing, we independently altered nucleotide identity and pairing at positions 12/79 and 15/76, two sites proximal to the DCL1 first cleavage site within the basal stem of the *shc* precursor and assessed their impact on amiRNA accumulation. For the 12/79 position, sRNA blot analysis showed higher accumulation of both amiRNAs from all modified precursor variants compared to wild-type configuration (Figure 3c). Notably, amiR-NbSu accumulation was significantly increased in samples expressing precursors with A12G or C79U modifications relative to the wild-type precursor, while amiR-NbDXS accumulated to significantly increased levels when expressed from variants with A12G or C79G. Similarly, at position 15/76, amiRNA accumulation was generally higher across all variants, with the U15G/U76C–U15C/U76G–U15G/U76 and U15G/U76C–U15C/U76G–U15A/U76–U15/U76A variants producing significantly higher amounts of amiR-NbSu and amiR-NbDXS, respectively (Figure 3d). Together, these results indicate that base pairing at internal positions 12/79 and 15/76 from *shc* basal stem generally increase amiRNA accumulation.

### Combined effects of multiple base-pairing modifications on amiRNA accumulation in *shc* precursors

To assess the cumulative impact of introducing multiple base-pair changes within the basal stem of the *shc* precursor, we generated double and triple mutants targeting positions 12/79, 15/76 and 18/73, regions located in close proximity to the DCL1 first cleavage site (Figure 4), and compared amiRNA accumulation from these variants to that observed in the corresponding single mutants. These modifications were selected based on previous analyses showing that single substitutions A12G, U15G and A18G significantly enhanced amiRNA accumulation (Figures 2–4). Given the comparable responses observed for amiR-NbSu and amiR-NbDXS in earlier experiments, only amiR-NbDXS-expressing constructs were used in this analysis for simplicity. All constructs were transiently expressed in *N. benthamiana* leaves, and amiRNA accumulation was analyzed at 2 dpa using sRNA blot assays.

**Figure 4.**
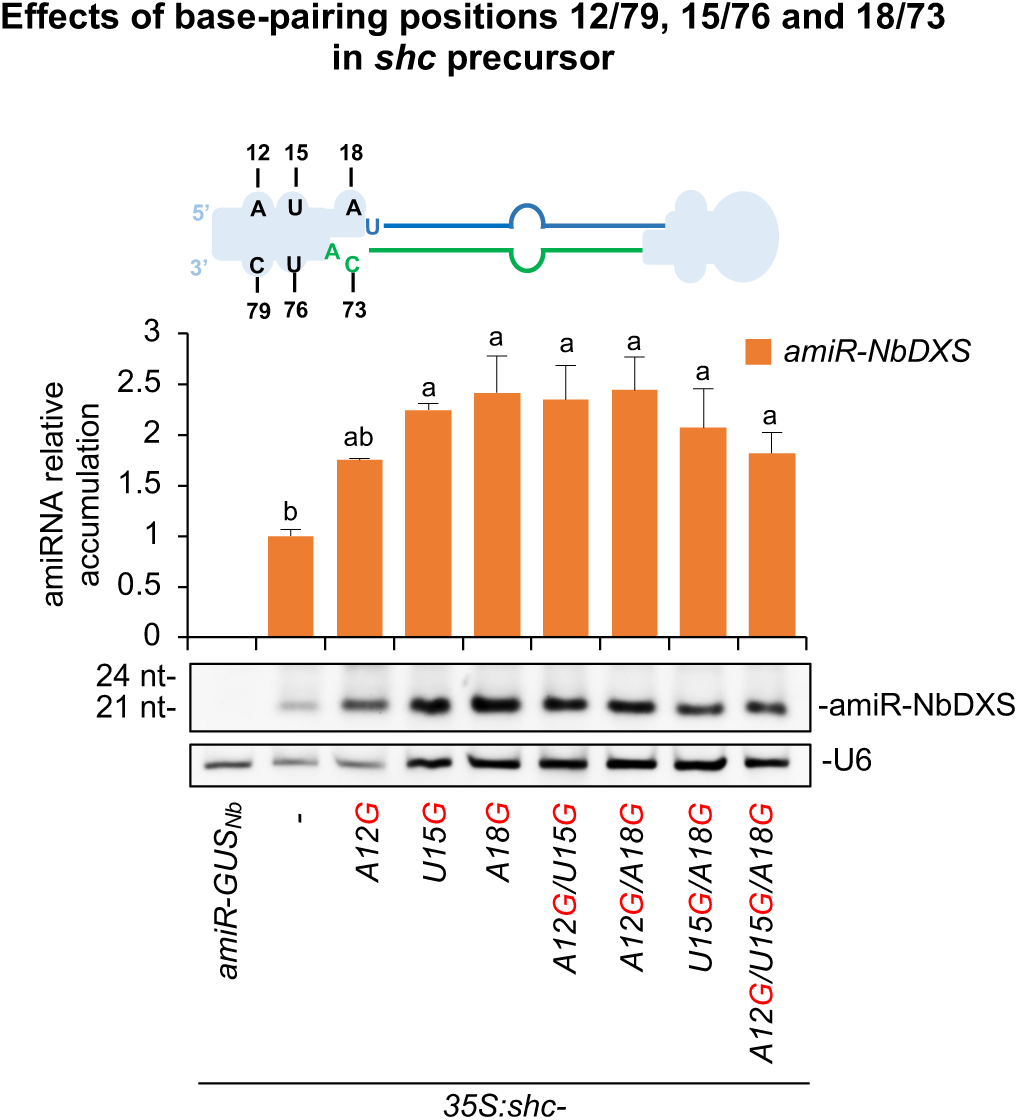
Functional analysis of wild-type and modified *shc-*based precursors without mismatches at positions 12/79, 15/76 and 18/73 in single, double and triple combinations. Top, diagram of wild-type *shc* amiRNA precursor with unpaired positions 12/79, 15/76 and 18/73 highlighted. Other details are as in Figure 2b. Bottom, Northern blot detection of amiR-NbDXS. Bars with the letter ‘a’ or ‘b’ are significantly different from that of the corresponding wild-type *shc-NbDXS or shc-A18G-NbDXS* samples, respectively (P < 0.05 in all pairwise Student’s *t*-test comparisons). Other details are as in Figure 2c.

All modified precursors produced significantly higher levels of amiR-NbDXS relative to the wild-type control (Figure 4). Among the single mutants, A18G supported the highest amiRNA accumulation; however, this increase was not significantly different from that observed with the U15G variant, while it was significantly higher than that conferred by the A12G variant, which showed reduced accumulation (Figure 4). Importantly, neither the double (A12G/U15G, A12G/A18G, U15G/A18G) nor the triple (A12G/U15G/A18G) mutants conferred further enhancement relative to the A18G single mutant, indicating a lack of additive or synergistic effects in this context. Based on these results, the *shc-A18G* variant, which reproducibly supports robust amiRNA accumulation, was selected for subsequent analyses.

### Processing accuracy of *shc* and *shc-A18G* precursors releasing amiR-NbDXS

To further confirm processing accuracy of *shc-A18G* precursors, sRNA libraries were prepared from plants expressing *35S:shc-A18G-NbDXS* and sequenced. For comparison, sRNA datasets from *35S:shc-NbDXS* samples were also analyzed (Cisneros *et al*., 2023). In both precursors, read coverage concentrated almost exclusively within the predicted amiRNA/amiRNA* region, with negligible accumulation along the remaining backbone (Figure 5a). The size profile was strongly dominated by 21-nt reads corresponding to the expected amiR-NbDXS sequence, while other sRNA species contributed only marginally. Moreover, reads were aligned relative to the amiR-NbDXS 5′ terminus, the high precision of processing became evident (Figure 5b). In both precursors, the vast majority of 21-nt reads, corresponding to authentic amiR-NbDXS, initiated exactly at position 0, with only minor offset reads detected at −1 or +1. The position-0 signal greatly exceeded neighboring positions for both precursors, reflecting the high 5′-end fidelity of the mature guide strand. Finally, quantification of processing accuracy, defined as the proportion of reads that perfectly matched the expected 21-nt amiR-NbDXS within the–4/+4 region surrounding the 5’ end, revealed a 90% accuracy for both precursors (Figure 5c). Collectively, these results show that the *shc-A18G* variant enhances amiR-NbDXS accumulation while fully preserving the defining features of accurate DCL1 processing: dominant 21-nt production, confinement of reads to the guide strand region, and a sharp, precisely defined 5’ end.

**Figure 5.**
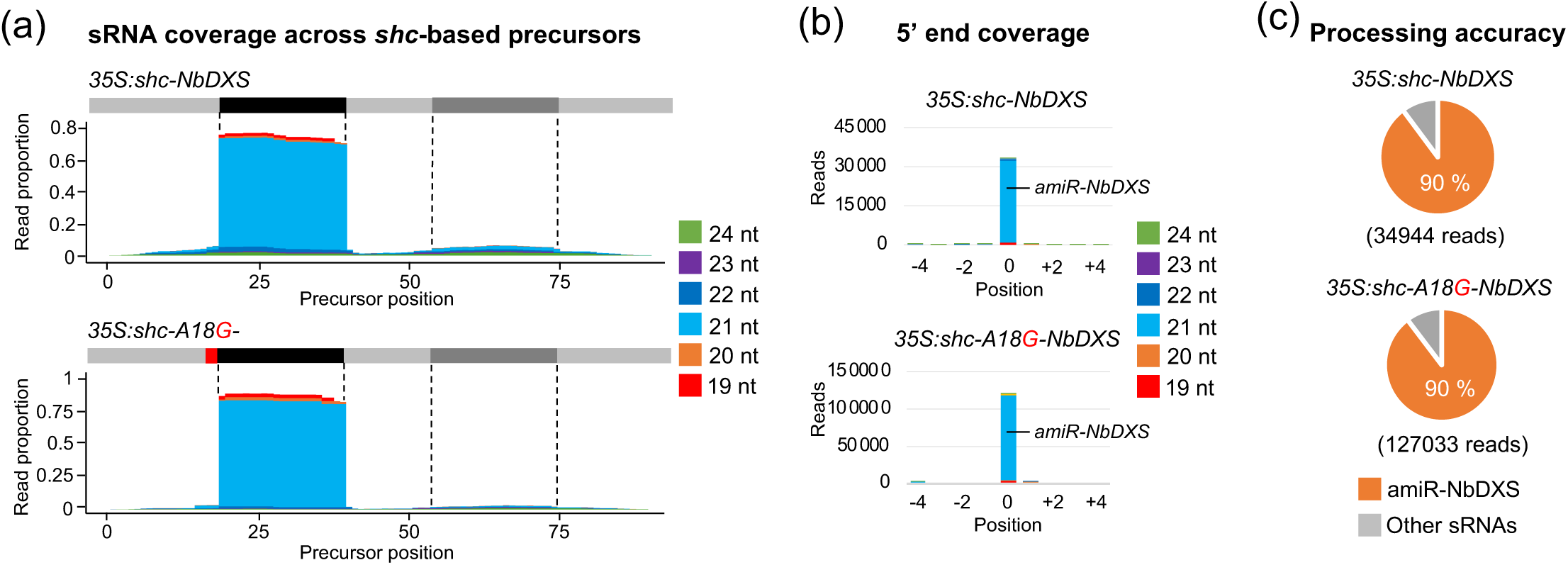
Processing of amiR-NbDXS from wild-type and A18G-modified *shc* precursors. (a) Small RNA (sRNA) coverage across *shc*-based precursors. The *x*-axis indicates the position on the precursor in nucleotides, from 5’ to 3’. At the top of each plot, the light gray line corresponds to the precursor backbone; the position of the amiRNA and amiRNA* in the precursors are indicated in black and dark gray respectively, and the A18G substitution in red. The *y*-axis is the sRNA coverage in proportion of reads for each nucleotidic position aligning to the positive strand. Coverage of reads of different lengths is shown in separate colors, stacked from bottom to top as indicated in the legend on the right. (b) sRNA 5’ coverage around the artificial miRNA (amiRNA) 5’ end in *shc*-based precursors. In the *x*-axis, 0 indicates the 5’ end of the amiRNA, -4 and +4 indicate 4 nt upstream and downstream of them. They *y*-axis is the sRNA 5’ coverage in total reads. The light blue portion of the bar at 0 represents authentic amiR-NbDXS reads. Other details are as in (a). (c) amiRNA processing accuracy from *shc*-based precursors. Pie charts show percentages of reads corresponding to expected, accurately processed 21-nt mature amiR-NbDXS (orange sectors) or to other 19-24-nt sRNAs (gray sectors).

### New high-throughput vectors for expressing amiRNAs from *shc-A18G*-based precursors

To facilitate high-throughput cloning and expression of amiRNAs from *shc-A18G* precursors, we developed two new “B/c” vectors incorporating the A18G-modified basal stem of *AtMIR390a* (Figure S1): (i) *pENTR-BS-AtMIR390a-A18G-B/c*, a Gateway-compatible entry vector enabling direct insertion of amiRNA sequences and subsequent recombination into preferred expression vectors with customizable promoters, terminators, and regulatory features; and (ii) *pMDC32B-BS-AtMIR390a-A18G-B/c*, a binary vector suitable for direct Agrobacterium-mediated transformation, eliminating intermediate subcloning steps (Figure S2). Both vectors contain the truncated *BS-AtMIR390a-A18G* region followed by a 1461-bp DNA cassette encoding the *ccd*B negative selection marker (Bernard and Couturier, 1992), flanked by two inverted *Bsa*I restriction sites positioned downstream of the precursor sequence. amiRNA constructs are generated using an established and cost-effective B/c cloning strategy (Cisneros *et al*., 2023; Carbonell *et al*., 2014). Briefly, amiRNA inserts are prepared by annealing two 58-nt overlapping and partially complementary oligonucleotides carrying the amiRNA sequence, with 5′-TGTG and 5′-AATG overhangs, and directionally ligated into *Bsa*I-digested *BS-AtMIR390a-A18G-B/c* vectors (Figure S2 and Text S1). These vectors were subsequently used throughout this study for functional validation of amiRNAs expressed from optimized *shc-A18G* precursors.

### Enhanced target silencing in transgenic Arabidopsis expressing A18G-modified *shc* amiRNA precursors

To evaluate the performance of the A18G-modified *shc* precursor in a stable genetic context, we generated transgenic Arabidopsis plants expressing amiRNAs amiR-AtFT, amiR-AtELF3 and amiR-AtCH42 (Figure 6a) targeting endogenous *FLOWERING LOCUS T* (*AtFT*), *EARLY FLOWERING 3* (*AtELF3*) or *CHLORINA 42* (*AtCH42*) endogenous genes, respectively, from either the wild-type or A18G-modified *shc* precursors. Efficient silencing of *AtFT*, *AtELF3* and *AtCH42* should result in significant delay in flowering time, hypocotyl elongation or to intense bleaching, respectively, as described before (Schwab *et al*., 2006; Kim and Somers, 2010). Briefly, we introduced amiR-AtFT, amiR-AtELF3 and amiR-AtCH42 into *pMDC32B-BS-AtMIR390a-A18G-B/c* to generate the *35S:shc-A18G-AtFT*, *35S:shc-A18G-AtELF3* and *35S:shc-A18G-AtCH42* constructs, respectively. These constructs were independently transformed into Arabidopsis Col-0 plants, along with control constructs *35S:shc-AtFT*, *35S:shc-AtELF3*, *35S:shc-AtCH42* and *35S:shc-GUS_At_*, which express an amiRNA targeting *GUS* (with no predicted off-targets in Arabidopsis) from the *shc* precursor (Cisneros *et al*., 2023). To systematically compare the processing and silencing efficacy of amiRNAs produced from wild-type versus A18G precursors, we analyzed plant phenotypes, amiRNA accumulation, target mRNA levels and processing accuracy in Arabidopsis T1 transgenic lines.

**Figure 6.**
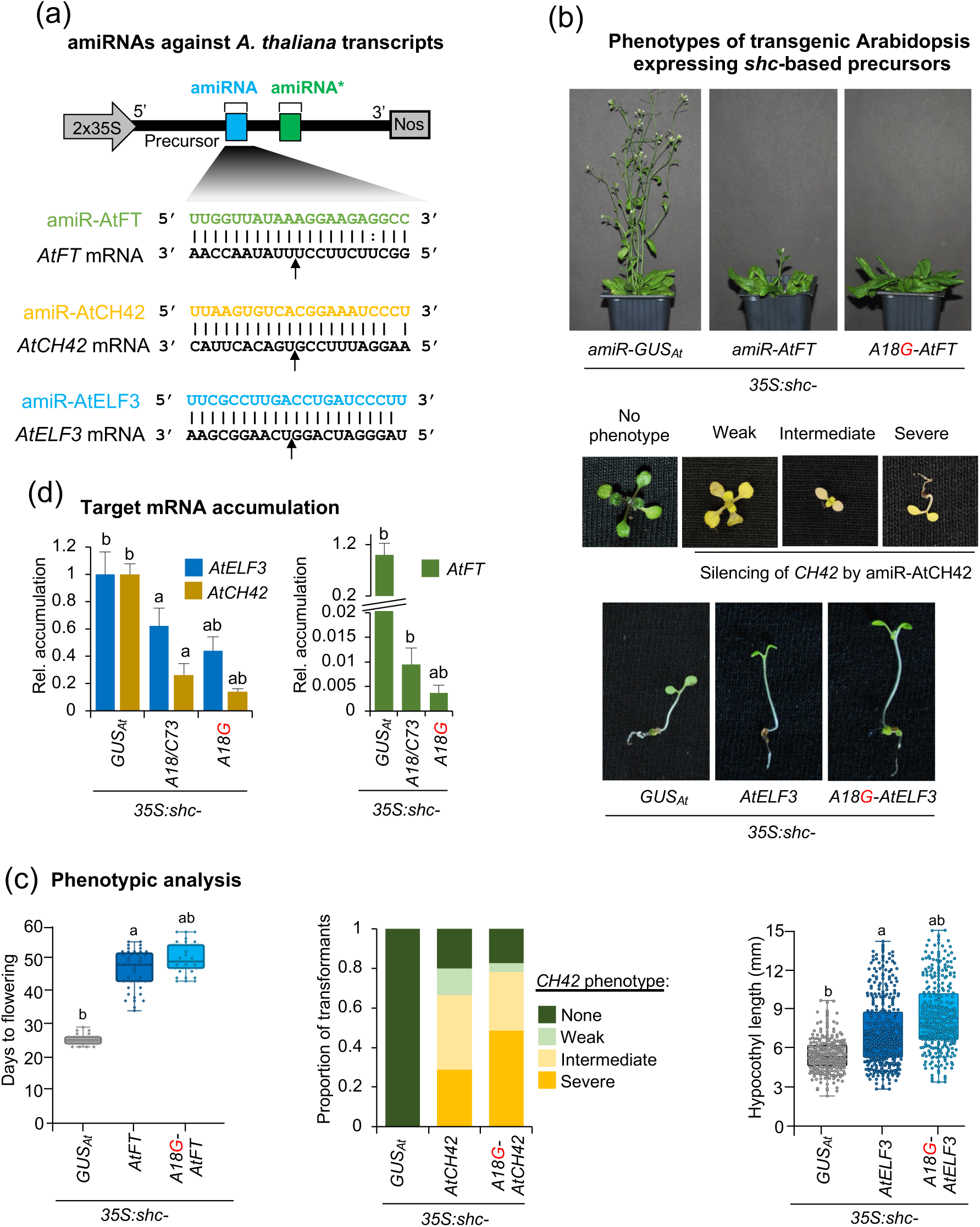
Functional analysis of constructs expressing the amiR-AtFT, amiR-AtCH42 and amiR-AtELF3 amiRNAs against Arabidopis *FLOWERING LOCUS T* (*AtFT*), *CHLORINE 42* (*AtCH42*) or *EARLY FLOWERING 3* (*AtELF3*) from wild-type and A18G-modified *shc* precursors. (a) Diagram of *shc-MIR390*-based amiRNA constructs including the base-pairing of amiRNAs and target mRNAs. Nucleotides corresponding to the guide strand of the amiRNA against *AtFT*, *AtCH42* and *AtELF3* are in green, yellow and blue, respectively, while nucleotides of target mRNAs are in black. The arrows indicate the amiRNA-predicted cleavage site. (b) Representative images of Arabidopsis T1 transgenic plants expressing amiRNAs from different precursors. Top, 45-day-old adult plants expressing amiR-GUS_At_ or amiR-AtFT. Middle, 10-day-old T1 seedlings expressing amiR-AtCH42 and showing bleaching phenotypes of diverse degrees. Bottom, 10-day-old T1 seedlings expressing amiR-AtELF3 and showing elongated hypocotyls. (c) Phenotyping analysis. Left, box plot representing the mean flowering time of Arabidopsis T1 transgenic plants expressing amiR-GUS_At_ or amiR-AtFT from different precursors. Center, bar graph representing, for each line, the proportion of seedlings displaying a severe (black areas), intermediate (dark gray areas), or weak (light gray areas) bleaching phenotype, or with wild-type appearance (white areas). Right, box plot representing the mean hypocotyl length of Arabidopsis T1 transgenic plants expressing amiR-GUS_At_ or amiR-AtELF3 from different precursors Pairwise Student’s t-test comparisons are represented with the letter ‘a’ or ‘b’ if significantly different from *35S:shc-GUS_At_*or wild-type *35S:shc-AtFT*/*35S:shc-AtELF3* samples, respectively (P < 0.05). (d) Target *AtFT*, *AtCH42* and *AtELF3* mRNA accumulation in RNA preparations from Arabidopsis plants [mean relative level (n = 3) + standard error] after normalization to ACTIN 2, as determined by quantitative RT-qPCR (*35S:shc-GUS_At_*= 1). Bars with the letter ‘a’ or ‘b’ are significantly different from *35S:shc-GUS_At_* or wild-type *35S:shc-AtFT*/*35S:shc-AtCH42/35S:shc-AtELF3* samples, respectively (P < 0.05).

Phenotypic analyses revealed that all *35S:shc-A18G-AtFT* (n = 34) transgenic lines expressing amiR-AtFT from the A18G-modified *shc* precursor exhibited a significantly delayed flowering time relative to those expressing the same amiRNA from the wild-type *shc* precursor (*35S:shc-AtFT*, n = 43), with mean flowering times of 46.6 ± 6 and 50 ± 4.7 days, respectively (Figure 6b–c, left; Table S1). Similarly, *35S:shc-A18G-AtCH42* seedlings expressing amiR-AtCH42 from the A18G-modified precursor displayed stronger bleaching phenotypes, with a higher proportion of individuals (48.7%) exhibiting severe chlorosis compared to those transformed with wild-type *shc 35S:shc-AtCH42* (28.8%) (Figure 6b–c, right; Table S1). In the case of amiR-AtELF3, phenotypic evaluation based on hypocotyl length under short-day conditions revealed that *35S:shc-A18G-AtELF3* transformants showed, on average, significantly enhanced hypocotyl elongation relative to *35S:shc-AtELF3* lines (Figure 6b–c, bottom; Table S1). These enhanced phenotypic effects were consistent with a stronger repression of their respective targets (Figure 6d) in lines expressing the A18G-modified precursor.

Finally, amiRNA accumulation and precursor processing was compared in Arabidopsis lines expressing amiR-AtFT, amiR-AtCH42 and amiR-ELF3 from wild-type or A18G-modified *shc* precursors (Figure 7). As shown by northern blot analysis, lines expressing amiRNAs from A18G-modified *shc* precursors accumulated significantly higher levels of amiRNAs, which migrated as single, discrete bands (Figure 7a). To analyze the accuracy of the processing and to confirm the presence of authentic amiRNAs, high throughput sequencing of sRNAs was performed from Arabidopsis lines expressing each amiRNA from wild-type or A18G-modifed *shc* precursors. Read coverage profiles revealed that sRNAs mapped almost exclusively to the predicted amiRNA/amiRNA* regions, with negligible reads along the remaining precursor backbone (Figure 7b). As with amiR-NbDXS-derived constructs, the distribution was strongly biased toward 21-nt sRNAs corresponding to the expected mature amiRNAs, with very limited contributions from other size classes. When reads were anchored to the amiRNA 5′ termini, both wild-type and A18G precursors showed a dominant peak at position 0, demonstrating highly precise DCL1 cleavage (Figure 7c). Importantly, processing accuracy was uniformly high across all amiRNAs, very similar for amiR-AtFT and amiR-AtELF3, and slightly higher for amiR-AtCH42 produced from *shc-A18G* precursors (Figure 7d).

**Figure 7.**
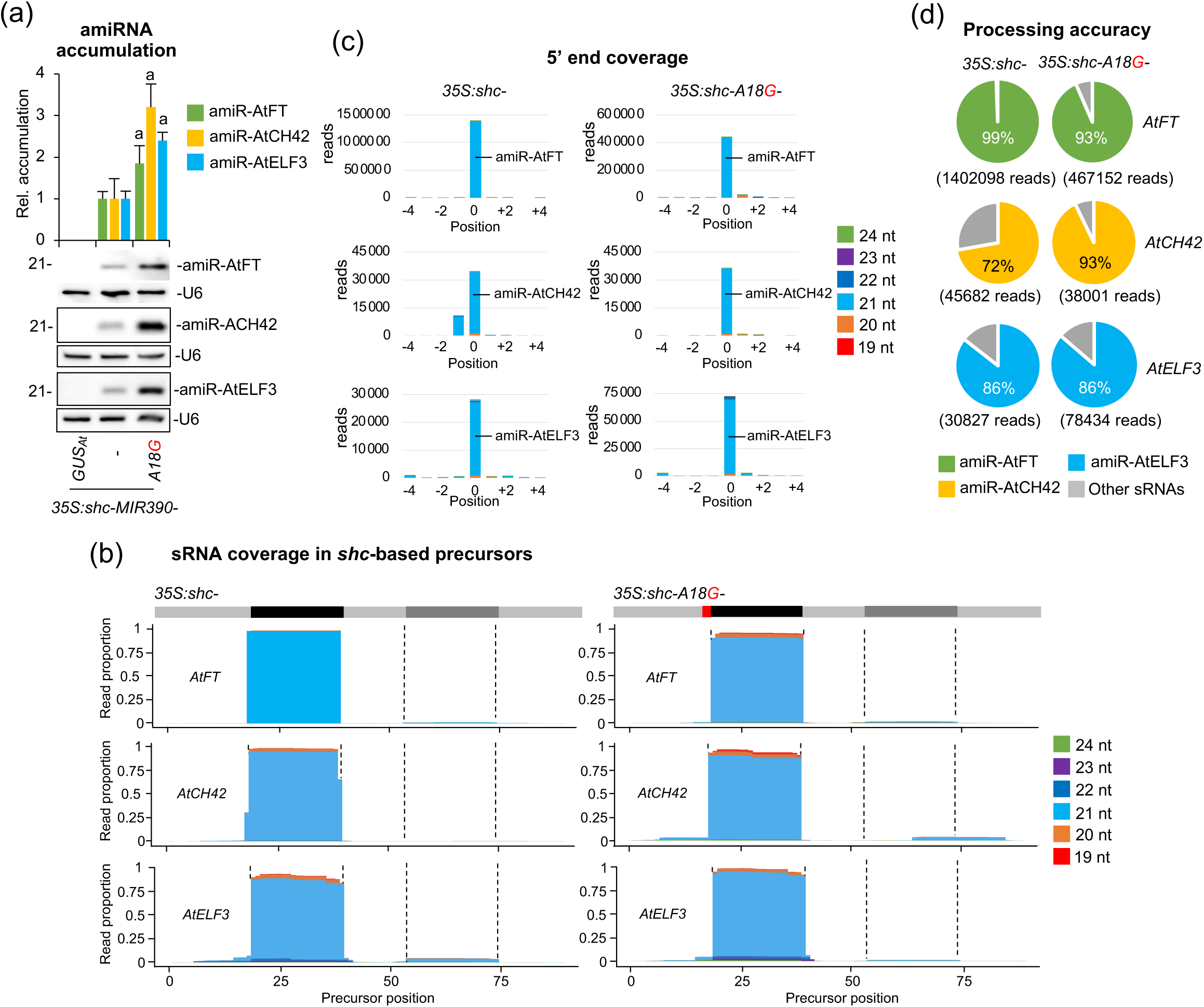
Accumulation and processing of amiR-AtFT, amiR-AtCH42 and amiR-AtELF3 amiRNAs from wild-type and A18G-modified *shc* precursors. (a) Northern blot detection of amiR-AtFT, amiR-AtCH42 and amiR-AtELF3 in RNA preparations from Arabidopsis plants. The graph at top shows the mean + standard deviation (n = 3) amiRNA relative accumulation (*35S:shc-MIR390-AtFT* = 1, *35S:shc-MIR390-AtCH42* = 1 and *35S:shc-MIR390-AtELF3* = 1). Bars with the letter ‘a’ are significantly different from that of *35S:shc-MIR390-AtFT , 35S:shc-MIR390-AtCH42* or *35S:shc-MIR390-AtELF3* control samples. One blot from three biological replicates is shown. Each biological replicate is a pool of at least nine independent lines selected randomly. U6 RNA blots are shown as loading controls. (b) Small RNA (sRNA) coverage across *shc*-based precursors. The *x*-axis indicates the position on the precursor in nucleotides, from 5’ to 3’. At the top of each plot, the light gray line corresponds to the precursor backbone; the position of the amiRNA and amiRNA* in the precursors are indicated in black and dark gray respectively, and the A18G substitution in red. The *y*-axis is the sRNA coverage in proportion of reads for each nucleotide position aligning to the positive strand. Coverage of reads of different lengths is shown in separate colors, stacked from bottom to top as indicated in the legend on the right. (c) sRNA 5’ coverage around the artificial miRNA (amiRNA) 5’ end in *shc*-based precursors. In the *x*-axis, 0 indicates the 5’ end of the amiRNA, -4 and +4 indicate 4 nt upstream and downstream of them. They *y*-axis is the sRNA 5’ coverage in total reads. The light blue portion of the bars at 0 represents authentic amiR-AtFT, amiR-AtCH42 and amiR-AtELF3 reads. Other details are as in (b). (d) amiRNA processing accuracy from *shc*-based precursors. Pie charts show percentages of reads corresponding to expected, accurately processed 21-nt mature amiR-AtFT, amiR-AtCH42 and amiR-AtELF3 (green, yellow and light blue sectors, respectively) or to other 19-24-nt sRNAs (gray sectors).

These analyses indicate that the A18G-modified precursor supports accurate and efficient DCL1 processing to release highly abundant amiRNAs. Altogether, these results show that stable expression of amiRNAs from the A18G-modified *shc* precursors produces increased levels of accurately processed amiRNAs for enhanced target silencing efficacy and specificity in transgenic Arabidopsis plants.

## DISCUSSION

In this study, we show that restoring base pairing at the nucleotide immediately upstream of the DCL1 first-cleavage site in both the native *AtMIR390a* and engineered *shc* precursors remarkably increases miRNA accumulation. These findings highlight the critical influence of precursor architecture on processing efficiency, and identify a novel *shc-A18G*-modified precursor with enhanced silencing activity across different species.

Systematic mutational analyses of several plant miRNA precursors revealed that efficient, high-fidelity miRNA biogenesis in plants depends on the structure of the precursor, particularly on the basal stem (Mateos *et al*., 2010; Song *et al*., 2010; Werner *et al*., 2010; Bajczyk *et al*., 2023; Li and Yu, 2021). In particular, earlier work has shown that the four positions flanking the miRNA/miRNA* duplex in natural Arabidopsis and eudicot miRNA precursors are usually base-paired and display position-specific sequence biases (Rojas *et al*., 2020). Interestingly, disrupting or restoring these pairings can substantially alter miRNA levels, with both the identity of the nucleotide and its precise position within the precursor affecting processing efficiency. For example, replacing the naturally mismatched 5’-U in *AtMIR172a* with a canonical C-G pair increased miR172a accumulation by 2.3 fold, whereas breaking adjacent pairs in *AtMIR172a* or *AtMIR164c* severely impaired processing (Mateos *et al*., 2010; Rojas *et al*., 2020). Conversely, introducing a mismatch at position 23 of *AtMIR164c* consistently reduced miRNA biogenesis (Rojas *et al*., 2020), and mutating the U-G pair at position 13/78 (U-G) in the basal stem of *AtMIR390a* significantly decreased miR390a levels (Cuperus, Montgomery, *et al*., 2010). Altogether, these findings support the idea that specific structural features of endogenous *MIRNA* precursors are critical for efficient processing. Our present data add position 18 of *AtMIR390a* to this catalogue, as pairing this nucleotide located immediately upstream to the mature miRNA substantially boosts miR390a accumulation. Remarkably, genome-wide analyses indicate that this site is generally paired across eudicots (Rojas *et al*., 2020), suggesting that the wild-type *AtMIR390a* architecture may have evolved not to maximize miR390a accumulation but to balance miR390a with *TAS3a* transcript levels. Such homeostasis is essential for proper accumulation of *TAS3a*-derived trans-acting siRNAs, which fine-tune auxin signaling and govern developmental processes such as leaf polarity and patterning, lateral-root formation, the timing of vegetative phase change and floral development (Fahlgren *et al*., 2006; Adenot *et al*., 2006; Marin *et al*., 2010; Garcia *et al*., 2006). Because A18G pairing enhances miR390a production (Figure 1b), we tested whether the same substitution would improve amiRNA biogenesis for targeted gene silencing. In both the full-length *AtMIR390a* and the recently described 89-nt *shc* minimal precursor (Cisneros *et al*., 2023), A18G significantly increased amiRNA levels. In addition, systematic mutagenesis of *shc* revealed that converting any of the three natural mismatches at positions 12, 15 and 18 into a G-C pairs significantly increased mature amiRNA levels, whereas pairing the second DCL1-cleavage site (U51) had no effect (Figures 2–3). The enhanced biogenesis likely reflects the higher thermodynamic stability of G–C pairs, which may rigidify the local stem and promote precise DCL1 activity. This interpretation aligns with genome-wide analyses showing that plant miRNA precursors are enriched for G–C/C–G pairs around the miRNA/miRNA* duplex (Rojas *et al*., 2020). Interestingly, combining two or three pairing mutations did not yield additive benefits (Figure 4), indicating that the *shc* precursor reaches a saturation point beyond which additional basal-stem stabilization no longer accelerates DCL1 processing. This plateau likely reflects the intrinsic rigidity of the chimera and suggests that optimal precursor recognition requires a balance between stem stability and the dynamic conformational changes mediated by HYL1 and SERRATE. Importantly, sRNA deep sequencing confirmed that DCL1 processing of the A18G-modified precursors is as accurate as that of the wild-type *shc* scaffold, with a very high proportion of reads correspond to the intended 21-nt amiRNA, with negligible alternative products (Figure 5 and 7). This high fidelity limits the release of ectopic sRNAs and therefore minimizes potential off-target effects. Moreover, no 21-nt secondary siRNAs in phase with the expected cleavage site were detected along the cognate target transcripts (Data S1), indicating that A18G-mediated silencing does not trigger RDR6-dependent transitivity and further reinforcing its specificity.

In conclusion, the *shc-A18G* backbone constitutes a minimal, high-efficiency platform for diverse gene-silencing applications. Its robust performance in *N. benthamiana* and Arabidopsis, across multiple guide sequences, confirms its broad utility. Moreover, the accompanying high-throughput B/c vectors, engineered with the A18G basal stem, simplify amiRNA construct assembly and cut oligonucleotide costs, an advantage for large-scale functional genomics screens (Jover-Gil *et al*., 2014; Hauser *et al*., 2013; Zhang *et al*., 2018), also reducing significantly the synthesis costs. Beyond basic research, this optimized amiRNA toolkit offers promise for agricultural, both in transgenic crops and in exogenous amiRNA treatments. Highly specific art-sRNA technologies represent an important step toward next-generation crops with improved resilience to environmental stresses and climate change.

## METHODS

### Plant species and growth conditions

*N. benthamiana* plants were cultivated in growth chambers maintained at 25 °C with a 12 h light/12 h dark photoperiod. *A. thaliana* ecotype Columbia-0 (Col-0) was grown at 22 °C under a 16 h light/8 h dark photoperiod, except for the *AtELF3* knock-down experiment, in which plants were plants were grown under a short-day regime of 8 h light/16 h dark photoperiod. Arabidopsis transformation was conducted via the floral dip method using *Agrobacterium tumefaciens* strain GV3101 as previously described (Clough and Bent, 1998). Selection and propagation of T1 transgenic lines followed standard protocols (López-Dolz *et al*., 2020). Plant images were captured using a Nikon D3000 digital camera equipped with an AF-S DX NIKKOR 18–55 mm f/3.5–5.6G VR lens.

### Arabidopsis phenotyping

Phenotypic analyses in *A. thaliana* were conducted in a blind manner as previously described (López-Dolz *et al*., 2020). Hypocotyl length was quantified from photographs of seedlings laid flat on agar plates alongside a ruler. Images were analyzed in ImageJ (Abramoff *et al*., 2004) by setting a scale based on the ruler, tracing hypocotyls using the segmented line tool, and extracting length values. Average hypocotyl lengths and standard deviations were calculated from these measurements. Flowering time was determined as the number of days from seed plating to the opening of the first floral bud (‘days to flowering’). A line was classified as exhibiting the ‘FT’ phenotype if its flowering time exceeded the average value observed in the *35S:shc-GUS_At_* control set. The ‘CH42’ phenotype was assessed in 10-day-old seedlings and categorized as ‘weak’, ‘intermediate’, or ‘severe’ based on the number of leaf primordia: more than two leaves (weak), exactly two leaves (intermediate), or no true leaves (severe; only cotyledons present). The ‘ELF3’ phenotype is scored in 10 days-old seedlings and was defined as a higher ‘hypocotyl’ value when compared to the average hypocotyl length value of the *35S:shc-GUS_At_* control set.

### Artificial small RNA design

P-SAMS script (https://github.com/carringtonlab/psams) (Fahlgren *et al*., 2016), configured to return unlimited optimal results, was used to obtain a list of optimal amiRNAs targeting *AtELF3* with high specificity (Data S2). Off-target filtering was applied using the *A. thaliana* transcriptome Araport 11 (https://ftp.ncbi.nlm.nih.gov/genomes/all/GCF/000/001/735/GCF_000001735.4_TAIR10.1/) (Cheng *et al*., 2017) to enhance amiRNA specificity. AmiR-GUS_Nb_, amiR-NbSu, amiR-NbDXS, amiR-GUS_At_, amiR-AtFT and amiR-AtCH42 guide sequences were described before (Cisneros *et al*., 2022; Schwab *et al*., 2006; López-Dolz *et al*., 2020).

### DNA constructs

Oligonucleotides AC-1268 and AC-1269 were annealed and ligated into *pENTR-D-TOPO* to generate *pENTR-BS-AtMIR390a-A18G-BB* including *AtMIR390a* basal stem sequence interrupted by two inverted *Bsa*I restriction sites. The *BS-AtMIR390a-A18G-BB* cassette from *pENTR-AtMIR390a-A18G-BB* was transferred by LR recombination into *pMDC32B* (Carbonell *et al*., 2014), a version of *pMDC32* (Curtis and Grossniklaus, 2003) with mutated *Bsa*I site, to generate *pMDC32B-BS-AtMIR390a-A18G-BB*. The B/c cassette was amplified from *pENTR-AtMIR390a-B/c* (Addgene plasmid #51778) with oligonucleotides AC-1270 and AC-1271, and ligated into *pENTR-D-TOPO*. Finally, the B/c cassette was excised by *Bsa*I digestion and inserted into *Bsa*I-digested *pENTR-BS-AtMIR390a-A18G-BB* and *pMDC32B-BS-AtMIR390a-A18G-BB* to generate *pENTR-BS-AtMIR390a-A18G-B/c* (Addgene plasmid 246715) and *pMDC32B-BS-AtMIR390a-A18G-B/c* (Addgene plasmid 246716) were deposited at Addgene (http://www.addgene.org/).

Constructs *35S:shc-GUS_Nb_*, *35S:AtMIR390a*, *35S:shc-C40G/U51C-NbSu*, *35S:shc-U51G-NbSu*, *35S:shc-C40A-NbSu*, *35S:shc-C40U/U51A-NbSu, 35S:shc-C40U/U51G-NbSu*, *35S:shc-C40G-NbSu, 35S:shc-C40G/U51C-NbDXS*, *35S:shc-U51G-NbDXS, 35S:shc-C40A-NbDXS*, *35S:shc-C40U/U51A-NbDXS*, *35S:shc-C40U/U51G-NbDXS*, *35S:shc-C40G-NbDXS*, *35S:shc-GUS_At_*, *35S:shc-AtELF3*, were obtained by ligating annealed oligonucleotide pairs AC-800/AC-801, AC-1272/AC-1273, AC-1114/AC-1115, AC-982/AC-983, AC-1116/AC-117, AC-1118/AC-1119, AC-1120/AC-1121, AC-1122/AC-1123, AC-1124/AC-1125, AC-886/AC-887, AC-1126/AC-1127, AC-1128/AC-1129, AC-1130/AC-1131, AC-1132/AC-1133, AC-1180/AC-1181, AC-1280/AC-1281, respectively, into *pMDC32B-BS-AtMIR390a-B/c* (Addgene plasmid #199560) (Cisneros *et al*., 2023).

Constructs *35S:AtMIR390a-A18G*, *35S:AtMIR390a-A18G-NbSu*, *35S:AtMIR390a-A18G-NbDXS*, *35S:shc-A18G-NbSu*, *35S:shc-C73U-NbSu*, *35S:shc-A18U/C73A-NbSu*, *35S:shc-A18U/C73G-NbSu*, *35S:shc-A18U/C73G-NbSu*, *35S:shc-A18G/C73U-NbSu*, *35S:shc-A18C/C73G-NbSu*, *35S:shc-A18G-NbDXS*, *35S:shc-A18C/C73G-NbDXS*, *35S:shc-C73U-NbDXS*, *35S:shc-A18U/C73A-NbDXS*, *35S:shc-A18U/C73G-NbDXS*, *35S:shc-A18G/C73U-NbDXS*, *35S:shc-A12G-_NbSu_*, *35S:shc-A12C/C79G-NbSu*, *35S:shc-C79U-NbSu*, *35S:shc-A12U/C79A-NbSu*, *35S:shc-A12U/C79G-NbSu*, *35S:shc-A12G/C79U-NbSu*, *35S:shc-A12G-NbDXS*, *35S:shc-A12C/C79G*, *35S:shc-C79U-NbDXS*, 3*5S:shc-A12U/C79A-NbDXS*, *35S:shc-A12U/C79G-NbDXS*, *35S:shc-A12G/C79U-NbDXS*, *35S:shc-U15G/U76C-NbSu*, *35S:shc-U15C/U76G-NbSu*, *35S:shc-U15A-NbSu*, *35S:shc-U76A-NbSu*, *35S:shc-U76G-NbSu*, *35S:shc-U15G-NbSu*, *35S:shc-U15G/U76C-NbDXS*, *35S:shc-U15C/U76G-NbDXS*, *35S:shc-U15A-NbDXS*, *35S:shc-U76A-NbDXS*, *35S:shc-U76G-NbDXS*, *35S:shc-U15G-NbDXS*, *35S:shc-A12G/U15G-NbDXS*, *35S:shc-A12G/A18G-NbDXS*, *35S:shc-U15G/A18G-NbDXS*, *35S:shc-A12G/U15G/A18G-NbDXS*, were obtained by ligating annealed oligonucleotide pairs AC-1274/AC-1275, AC-1286/AC-1287, AC-1288/AC-1289, AC-949/AC-950, AC-951/AC-952, AC-953/AC-954, AC-955/AC-956, AC-957/AC-958, AC-959/AC-960, AC-961/AC-962, AC-878/AC-879, AC-969/AC-970, AC-963/AC-964, AC-965/AC-966, AC-967/AC-968, AC-974/AC-975, AC-1150/AC-1151, AC-1152/AC-1153, AC-1154/AC-1155, AC-1158/AC-1159, AC-1156/AC-1157, AC-882/AC-883, AC-1160/AC-1161, AC-1162/AC-1163, AC-1164/AC-1165, AC-1168/AC-1169, AC-1166/AC-1167, AC-1077/AC-1078, AC-1134/AC-1135, AC-976/AC-977, AC-1136/AC-1137, AC-1140/AC-1141, AC-1138/AC-1139, AC-1085/AC-1086, AC-1142/AC-1143, AC-884/AC-885, AC-1144/AC-1145, AC-1148/AC-1149, AC-1146/AC-1147, AC-1237/AC-1238, AC-1087/AC-1088, AC-1239/AC-1240, AC-1241/AC-1242, respectively, into *pMDC32B-B/c* (Addgene plasmid #227963) (Cisneros *et al*., 2025).

Constructs *35S:shc-A18G-AtFT*, *35S:shc-A18G-AtCH42* and 35S*:shc-A18G-AtELF3* were obtained by ligating annealed oligonucleotide pairs AC-1276/AC-1277, AC-1278/AC-1279 and AC-1281/AC-1282, respectively, into *pMDC32B-BS-AtMIR390a-A18G-B/c* (Addgene plasmid # 246716). A detailed protocol for cloning amiRNAs in in new B/c vectors is described in Text S1. Constructs *35S:GUS*, *35S:AtMIR390a-GUS_Nb_*, *35S:AtMIR390a-NbSu*, *35S:AtMIR390a-NbDXS*, *35S:shc-NbSu*, *35S:shc-NbDXS*, *35S:shc-AtFT* and *35S:shc-AtCH42* were described before (Montgomery *et al*., 2008; Cisneros *et al*., 2022; Cisneros *et al*., 2023). The sequences of all miRNA/amiRNA precursors are listed in Text S2. The sequences of newly developed B/c vectors are listed in Text S3.

### Transient expression of constructs

Agrobacterium-mediated infiltration of constructs into *N. benthamiana* leaves was performed as previously described (Carbonell *et al*., 2015; Cuperus, Carbonell, *et al*., 2010).

### Chlorophyll extraction and analysis

Chlorophyll and other pigments were extracted from *N. benthamiana* leaves and analyzed as previously described (López-Dolz *et al*., 2020; Carbonell *et al*., 2015).

### Total RNA preparation

Total RNA from *N. benthamiana* leaves or Arabidopsis seedlings or inflorescences was isolated as previously described (Cisneros *et al*., 2023). Each sample consisted of pools of two *N. benthamiana* leaves or 9–12 Arabidopsis seedlings or inflorescences, prepared in triplicate.

### Real-time RT-qPCR

Real-time RT-qPCR was performed using the RNA samples previously analyzed by sRNA blotting as described (Cisneros *et al*., 2025). Primer sequences are listed in Table S2. Target mRNA expression was quantified relative to the reference gene *NbPP2A* or *AtACT2* in *N. benthamiana* and Arabidopsis, respectively, using the ΔΔCt comparative method in QuantStudio Design and Analysis software (v1.5.1; Thermo Fisher Scientific). Three independent biological replicates, each with two technical replicates, were analyzed.

### Small RNA blot assays

Total RNA (20 µg) was separated on 17% polyacrylamide gels (0.5× TBE, 7 M urea) and transferred onto positively charged nylon membranes. DNA or LNA probes were labeled using the second-generation DIG Oligonucleotide 3’-End Labeling Kit (Roche), and hybridizations were performed at 38°C as described (Tomassi *et al*., 2017). Membranes were imaged using an ImageQuant 800 CCD imager (Cytiva), and signals were quantified with ImageQuant TL software (v10.2; Cytiva). Probe sequences are listed in Table S2.

### Small RNA sequencing and data analysis

Total RNA quality, purity, and integrity were verified using an Agilent 2100 Bioanalyzer (RNA 6000 Nano kit) prior to sRNA library preparation and single-end 50-nt sequencing (SE50) performed by BGI (Hong Kong, China) on a DNBSEQ-G400 sequencer. Adapter-trimmed and quality-filtered reads provided by BGI were collapsed using the FASTX-Toolkit (http://hannonlab.cshl.edu/fastx_toolkit) to merge identical sequences while preserving read counts. Each unique read was mapped to the forward strand of the corresponding amiRNA precursor (Data S3) using a custom Python script deposited at Github (https://github.com/acarbonell/map_sRNA_reads/) that allowed exact matches without gaps or mismatches, calculating read counts and reads per million mapped reads (RPM). sRNA alignments were visualized using sRNA_Viewer software (Axtell Lab, Pennsylvania State University; https://github.com/MikeAxtell/sRNA_Viewer). Processing accuracy was assessed by quantifying the proportion of 19–24 nt sRNA (+) reads mapping within ±4 nt of the predicted amiRNA guide’s 5′ end (Carbonell *et al*., 2015; Cuperus, Carbonell, *et al*., 2010).

### Statistical analysis

Statistical analyses are detailed in the figure legends. Significant differences were determined using a two-tailed Student’s *t*-test.

## ACCESSION NUMBERS

Gene identifiers used in this study are as follows. Arabidopsis: *AtACT2* (AT3G18780), *AtCH42* (AT4G18480), *AtELF3* (AT2G25930) and *AtFT* (AT1G65480). *N. benthamiana: NbSu* (Nbv5.1tr6204879), *NbDXS* (Nbv5.1tr6224823), *NbPP2A* (Nbv5.1tr6224808). The *Escherichia coli* β-glucuronidase (GUS) gene sequence corresponds to GenBank accession S69414.1. High-throughput sequencing data can be found in the Sequence Read Archive (SRA) database under accession number PRJNA957136 and PRJNA1312446.

## ACKOWLEDGEMENTS

We thank IBMCP greenhouse staff for their invaluable help maintaining the plants. This work was supported by grants or fellowships from MICIU/AEI/10.13039/501100011033/FEDER, UE [PID2024-155602OB-100 and PID2021-122186OB-100 to A.C.; PRE2022-102565 to J.L.G], from Consejo Superior de Investigaciones Científicas (CSIC, Spain) [JAEINT_22_02953 and JAEINT_ 23_01968 to P.G.C. and S.R.R., respectively].

## AUTHOR CONTRIBUTIONS

J-J.L.G. did the majority of the experimental work with assistance from of P.G.C., S.R.R., L.D.C. and S.T.F. J-J.L.G. and A.C. analyzed the data. A.C. conceived the study, supervised the project and wrote the manuscript with input from all authors.

## SUPPORTING MATERIAL

**Data S1.** 21-nt sRNA reads mapping amiRNA targets in amiRNA-expressing tissues.

**Data S2**. P-SAMS designs of art-sRNA sequences.

**Data S3**. sRNA reads mapping amiRNA precursors in amiRNA-expressing tissues.

**Figure S1.** *BS-AtMIR390a-A18G-B/c*-based vectors for direct cloning of amiRNAs.

**Figure S2.** Direct amiRNA cloning in *AtMIR390a-A18G-B/c* (*Bsa*I/*ccd*B)-based vectors including a *ccd*B cassette flanked by two *Bsa*I sites.

**Table S1**. Phenotypic penetrance of amiRNAs expressed in *A. thaliana* Col-0 T1 transgenic.

**Table S2**. Name, sequence and use of oligonucleotides used in this study.

**Text S1**. Protocol to design and clone amiRNAs in *BS-AtMIR390a-A18G-B/c*-based vectors *pENTR-BS-AtMIR390a-A18G-B/c* and *pMDC32B-BS-AtMIR390a-A18G-B/c*.

**Text S2**. DNA sequence in FASTA format of all precursors used to express amiRNAs in plants

**Text S3**. DNA sequence of *BS-AtMIR390a-A18G-B/c*-based vectors used for direct cloning of amiRNAs.

