## Supporting_Figures_Tables_Texts for "Enhanced MicroRNA Accumulation and Gene Silencing Efficiency through Optimized Precursor Base Pairing"

#### SUPPORTING MATERIAL

**Data S1.** 21-nt sRNA reads mapping amiRNA targets in amiRNA-expressing tissues.

**Data S2**. P-SAMS designs of art-sRNA sequences.

**Data S3**. sRNA reads mapping amiRNA precursors in amiRNA-expressing tissues.

**Figure S1**. *BS-AtMIR390a-A18G-B/c*-based vectors for direct cloning of amiRNAs.

**Figure S2**. Direct amiRNA cloning in *AtMIR390a-A18G-B/c* (*Bsa*I/*ccd*B)-based vectors including a *ccd*B cassette flanked by two *Bsa*I sites.

**Table S1**. Phenotypic penetrance of amiRNAs expressed in *A. thaliana* Col-0 T1 transgenic.

**Table S2**. Name, sequence and use of oligonucleotides used in this study.

**Text S1**. Protocol to design and clone amiRNAs in *BS-AtMIR390a-A18G-B/c*-based vectors *pENTR-BS-AtMIR390a-A18G-B/c* and *pMDC32B-BS-AtMIR390a-A18G-B/c*.

**Text S2**. DNA sequence in FASTA format of all precursors used to express amiRNAs in plants.

**Text S3**. DNA sequence of *BS-AtMIR390a-A18G-B/c*-based vectors used for direct cloning of amiRNAs.

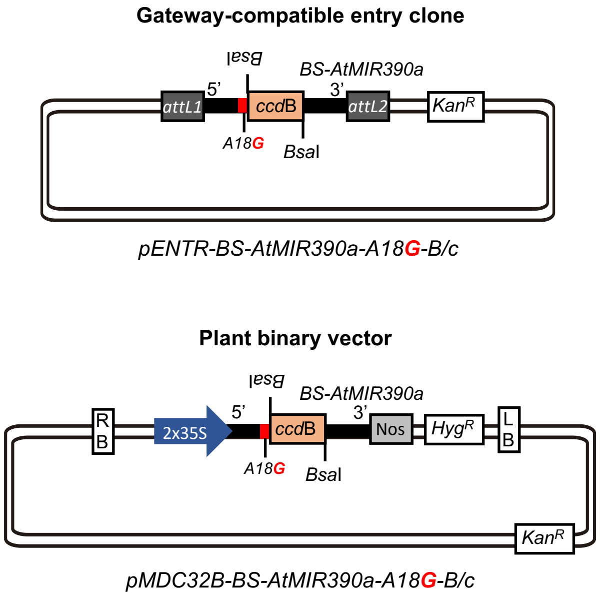


**Figure S1**. *BS-AtMIR390a-A18G-B/c*-based vectors for direct cloning of amiRNAs. Top, diagram of the Gateway-compatible *pENTR-BS-AtMIR390a-A18G-B/c* entry vector. Bottom, diagram of the *pMDC32B-BS-AtMIR390a-B/c* binary vector for in plant expression of amiRNAs. RB: right border; BS, basal stem; 35S: Cauliflower mosaic virus promoter; *Bsa*I: *Bsa*I recognition site, *ccd*B: gene encoding the gyrase toxin; LB: left border; attL1 and attL2: GATEWAY recombination sites. *Kan*$^{R}$: kanamycin resistance gene; *Hyg*$^{R}$: hygromycin resistance gene. The A18G mutation at the 5' BS is shown in red.

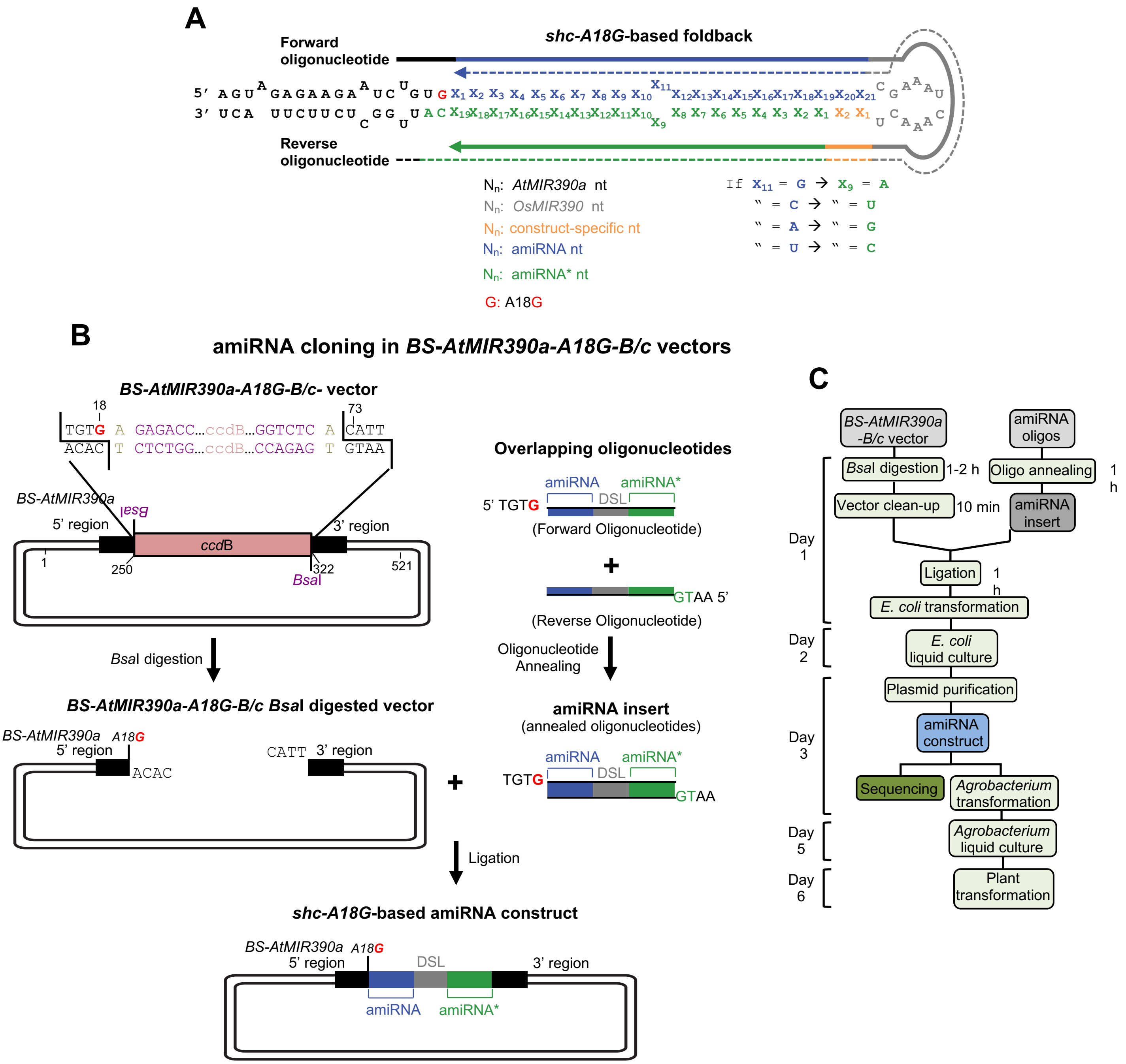


**Figure S2.** Direct cloning of amiRNAs in vectors containing a modified version of *BS-AtMIR390a-A18G* that includes a *ccd*B cassette flanked by two *Bsa*I sites (*Bsa*I/*ccd*B or ‘B/c’ vectors). A, Design of two overlapping oligonucleotides for amiRNA cloning in *BS-AtMIR390a-A18G*-based ”B/c” vectors including *OsMIR390* DSL sequences to produce a final *shc* precursor. Sequences covered by the forward and the reverse oligonucleotides are represented with continuous or dotted lines, respectively. Nucleotides of *BS-AtMIR390a-A18G* precursor, *OsMIR390*-derived distal stem loop (DSL), amiRNA guide strand and amiRNA* strand are in black, grey, blue and green, respectively, except A18G which is in red. Other nucleotides that may be modified for preserving authentic *OsMIR390a* foldback secondary structure are in orange. Rules for assigning identity to position 9 of the amiRNA* are indicated. B, Diagram of the steps for amiRNA cloning in *BS-AtMIR390a-A18G-B/c* vectors. The amiRNA insert obtained after annealing the two overlapping oligonucleotides has 5'-TGTG and 5'-AATG overhangs and is directly inserted in a directional manner into a *BS-AtMIR390a-A18G-B/c* vector previously linearized with *Bsa*I. Nucleotides of the *Bsa*I sites and those arbitrarily chosen and used as spacers between the *Bsa*I recognition sites and the *BS-AtMIR390a-A18G* sequence are in purple and light brown, respectively. Other details are as described in panel A. C, Flowchart of steps from amiRNA construct generation to plant transformation.

**Table S1:** Phenotypic penetrance of amiRNAs expressed in *A. thaliana* Col-0 T1 transgenic plants

| **Construct** | **T1 analyzed** | **Phenotypic penetrance[a]** |
|---|---|---|
| *35S:shc-amiR-GUS$_{At}$* | 44 | 0% |
| *35S:shc-amiR-AtFT* | 43 | 100% |
| *35S:shc-A18G-amiR-AtFT* | 34 | 100% |
| *35S:shc-amiR-GUS$_{At}$* | 278 | 0% |
| *35S:shc-amiR-AtELF3* | 398 | 72.4% |
| *35S:shc-A18G-amiR-AtELF3* | 335 | 89% |
| *35S:shc-amiR-GUS$_{At}$* | 260 | 0% |
| *35S:shc-amiR-AtCH42* | 125 | 100%<br>13.6% weak<br>37.6% intermediate<br>28.8 % severe |
| *35S:shc-A18G-amiR-AtCH42* | 230 | 100%<br>4.3% weak<br>29.6% intermediate<br>48.7 % severe |

[a] The FT phenotype was defined as a higher 'days to flowering' value when compared to the average 'days to flowering' value of the *35S:pri-amiR-GUS$_{Ath}$* control set. The ELF3 phenotype is scored in 10 days-old seedlings and was defined as a higher 'hypocotyl' value when compared to the average hypocotyl length value of the *35S:shc-amiR-GUS$_{At}$* control set. CH42 phenotype is scored in 10 days-old seedling and is considered 'weak', 'intermediate' or 'severe' if seedlings have >2 leaves, exactly two leaves or no leaves (only two cotyledons), respectively.

**Table S2.** Name, sequence and use of DNA oligonucleotides used in this study.

| Name | Sequence | Type* | Construct/Aim |
|---|---|---|---|
| AC-55 | AGGGGCCATGCTAATCTTCTC | ssDNA | Probe for U6 detection |
| AC-157 | GGCCTCTTCCTTTATAACCAA | ssDNA | Probe for amiR-AtFT detection |
| AC-158 | AGGGATTTCCGTGACACTTAA | ssDNA | Probe for amiR-AtCH42 detection |
| AC-159 | AAAAATGGCTGAGGCTGATGA | ssDNA | qPCR amplification of *AtACT2* mRNA |
| AC-160 | GAAAAACAGCCCTGGGAGC | ssDNA | |
| AC-163 | CATGCACAAGTAGGGACGGTT | ssDNA | qPCR amplification of *AtCH42* mRNA |
| AC-164 | GTCACGGAAATCCTTTGGGTT | ssDNA | |
| AC-169 | TGGAACAACCTTTGGCAATG | ssDNA | qPCR amplification of *AtFT* mRNA |
| AC-170 | CGACACGATGAATTCCTGCA | ssDNA | |
| AC-355 | GACCCTGATGTTGATGTTCGCT | ssDNA | qPCR amplification of *NbSu* mRNA |
| AC-356 | GAGGGATTTGAAGAGAGATTTC | ssDNA | |
| AC-359 | GGTGGTGGGACTGGTATGAA | ssDNA | qPCR amplification of *NbDXS* mRNA |
| AC-360 | GCAAATCTCACTGGCAGCTT | ssDNA | |
| AC-365 | GACCCTGATGTTGATGTTCGCT | ssDNA | qPCR amplification of *NbPP2A* mRNA |
| AC-366 | GAGGGATTTGAAGAGAGATTTC | ssDNA | |
| AC-417 | G+CGG+GAA+GTC+CAC+CAC+GGT+TA | ssLNA | Probe for amiR-NbSu detection |
| AC-418 | C+TGT+TAG+GAA+CCC+GCG+GTT+TA | ssLNA | Probe for amiR-NbDXS detection |
| AC-587 | AAGGGATCAGGTCAAGGCGAA | ssDNA | Probe for amiR-AtPDS3 detection |
| AC-591 | ATTGCTGCATCACCGGATCT | ssDNA | qPCR amplification of *AtELF3* |
| AC-592 | TCACCCCTTTGTTTGACGACA | ssDNA | |
| AC-800 | TGTATCTTGTAACGCGCTTTCCCAGCGAAATCAAACTCT GGGAAAGCTCGTTACAAGA | ssDNA | Generation of *35S:shc-GUS*$_{Nb}$ |
| AC-801 | AATGTCTTGTAACGAGCTTTCCCAGAGTTTGATTTCGCTG GGAAAGCGCGTTACAAGA | ssDNA | |
| AC-878 | TGTAAGTAGAGAAGAATCTGTGTAAACCGCGGGTTCCTA ACAGCGAAATCAAACTCTGTTAGGAAACCGCGGTTTACA TTGGCTCTTCTTACT | ssDNA | Generation of *35S:shc-A18G-NbDXS* |
| AC-879 | AATGAGTAAGAAGAGCCAATGTAAACCGCGGTTTCCTA ACAGAGTTTGATTTCGCTGTTAGGAACCCGCGGTTTACA CAGATTCTTCTCTACT | ssDNA | |
| AC-882 | TGTAAGTAGAGAAGAGTCTGTATAAACCGCGGGTTCCTA ACAGCGAAATCAAACTCTGTTAGGAAACCGCGGTTTACA TTGGCTCTTCTTACT | ssDNA | Generation of *35S:shc-A12G-NbDXS* |
| AC-883 | AATGAGTAAGAAGAGCCAATGTAAACCGCGGTTTCCTA ACAGAGTTTGATTTCGCTGTTAGGAACCCGCGGTTTATA CAGACTCTTCTCTACT | ssDNA | |
| AC-884 | TGTAAGTAGAGAAGAATCAGTATAAACCGCGGGTTCCT AACAGCGAAATCAAACTCTGTTAGGAAACCGCGGTTTAC ATTGGCTCTTCTTACT | ssDNA | Generation of *35S:shc-U15A-NbDXS* |
| AC-885 | AATGAGTAAGAAGAGCCAATGTAAACCGCGGTTTCCTA ACAGAGTTTGATTTCGCTGTTAGGAACCCGCGGTTTATA CTGATTCTTCTCTACT | ssDNA | |
| AC-886 | TGTATAAACCGCGGGTTCCTAACAGCGAAATCAAACGCT GTTAGGAAACCGCGGTTTA | ssDNA | Generation of *35S:shc-U51G-NbDXS* |
| AC-887 | AATGTAAACCGCGGTTTCCTAACAGCGTTTGATTTCGCT GTTAGGAACCCGCGGTTTA | ssDNA | |
| AC-949 | TGTAAGTAGAGAAGAATCTGTGTAACCGTGGTGGACTTC CCGCCGAAATCAAACTGCGGGAAGTCAACCACGGTTAC ATTGGCTCTTCTTACT | ssDNA | Generation of *35S:shc-A18G-NbSu* |
| AC-950 | AATGAGTAAGAAGAGCCAATGTAACCGTGGTTGACTTCC CGCAGTTTGATTTCGGCGGGAAGTCCACCACGGTTACAC AGATTCTTCTCTACT | ssDNA | |
| AC-951 | TGTAAGTAGAGAAGAATCTGTATAACCGTGGTGGACTTC CCGCCGAAATCAAACTGCGGGAAGTCAACCACGGTTAT ATTGGCTCTTCTTACT | ssDNA | Generation of *35S:shc-C73U-NbSu* |

| | | | |
|---|---|---|---|
| AC-952 | AATGAGTAAGAAGAGCCAATATAACCGTGGTTGACTTCCCGCAGTTTGATTTCGGCGGGAAGTCCACCACGGTTATACAGATTCTTCTCTACT | ssDNA | |
| AC-953 | TGTAAGTAGAGAAGAATCTGTTTAACCGTGGTGGACTTCCCGCCGAAATCAAACTGCGGGAAGTCAACCACGGTTAAATTGGCTCTTCTTACT | ssDNA | Generation of *35S:shc-A18U/C73A-NbSu* |
| AC-954 | AATGAGTAAGAAGAGCCAATTTAACCGTGGTTGACTTCCCGCAGTTTGATTTCGGCGGGAAGTCCACCACGGTTAAACAGATTCTTCTCTACT | ssDNA | |
| AC-955 | TGTAAGTAGAGAAGAATCTGTTTAACCGTGGTGGACTTCCCGCCGAAATCAAACTGCGGGAAGTCAACCACGGTTAGATTGGCTCTTCTTACT | ssDNA | Generation of *35S:shc-A18U/C73G-NbSu* |
| AC-956 | AATGAGTAAGAAGAGCCAATCTAACCGTGGTTGACTTCCCGCAGTTTGATTTCGGCGGGAAGTCCACCACGGTTAAACAGATTCTTCTCTACT | ssDNA | |
| AC-957 | TGTAAGTAGAGAAGAATCTGTGTAACCGTGGTGGACTTCCCGCCGAAATCAAACTGCGGGAAGTCAACCACGGTTATATTGGCTCTTCTTACT | ssDNA | Generation of *35S:shc-A18G/C73U-NbSu* |
| AC-958 | AATGAGTAAGAAGAGCCAATATAACCGTGGTTGACTTCCCGCAGTTTGATTTCGGCGGGAAGTCCACCACGGTTACACAGATTCTTCTCTACT | ssDNA | |
| AC-959 | TGTAAGTAGAGAAGAATCTGTCTAACCGTGGTGGACTTCCCGCCGAAATCAAACTGCGGGAAGTCAACCACGGTTAGATTGGCTCTTCTTACT | ssDNA | Generation of *35S:shc-A18C/C73G-NbSu* |
| AC-960 | AATGAGTAAGAAGAGCCAATCTAACCGTGGTTGACTTCCCGCAGTTTGATTTCGGCGGGAAGTCCACCACGGTTAGACAGATTCTTCTCTACT | ssDNA | |
| AC-961 | TGTAAGTAGAGAAGAATCTGTATAAACCGCGGGTTCCTAACAGCGAAATCAAACTCTGTTAGGAAACCGCGGTTTATATTGGCTCTTCTTACT | ssDNA | Generation of *35S:shc-C73U-NbDXS* |
| AC-962 | AATGAGTAAGAAGAGCCAATATAAACCGCGGTTTCCTAACAGAGTTTGATTTCGCTGTTAGGAACCCGCGGTTTATACAGATTCTTCTCTACT | ssDNA | |
| AC-963 | TGTAAGTAGAGAAGAATCTGTTTAAACCGCGGGTTCCTAACAGCGAAATCAAACTCTGTTAGGAAACCGCGGTTTAAATTGGCTCTTCTTACT | ssDNA | Generation of *35S:shc-A18U/C73A-NbDXS* |
| AC-964 | AATGAGTAAGAAGAGCCAATTTAAACCGCGGTTTCCTAACAGAGTTTGATTTCGCTGTTAGGAACCCGCGGTTTAAACAGATTCTTCTCTACT | ssDNA | |
| AC-965 | TGTAAGTAGAGAAGAATCTGTTTAAACCGCGGGTTCCTAACAGCGAAATCAAACTCTGTTAGGAAACCGCGGTTTAGATTGGCTCTTCTTACT | ssDNA | Generation of *35S:shc-A18U/C73G-NbDXS* |
| AC-966 | AATGAGTAAGAAGAGCCAATCTAAACCGCGGTTTCCTAACAGAGTTTGATTTCGCTGTTAGGAACCCGCGGTTTAAACAGATTCTTCTCTACT | ssDNA | |
| AC-967 | TGTAAGTAGAGAAGAATCTGTGTAAACCGCGGGTTCCTAACAGCGAAATCAAACTCTGTTAGGAAACCGCGGTTTATATTGGCTCTTCTTACT | ssDNA | Generation of *35S:shc-A18G/C73U-NbDXS* |
| AC-968 | AATGAGTAAGAAGAGCCAATATAAACCGCGGTTTCCTAACAGAGTTTGATTTCGCTGTTAGGAACCCGCGGTTTACACAGATTCTTCTCTACT | ssDNA | |
| AC-969 | TGTAAGTAGAGAAGAATCTGTCTAAACCGCGGGTTCCTAACAGCGAAATCAAACTCTGTTAGGAAACCGCGGTTTAGATTGGCTCTTCTTACT | ssDNA | Generation of *35S:shc-A18C/C73G-NbDXS* |
| AC-970 | AATGAGTAAGAAGAGCCAATCTAAACCGCGGTTTCCTAACAGAGTTTGATTTCGCTGTTAGGAACCCGCGGTTTAGACAGATTCTTCTCTACT | ssDNA | |
| AC-974 | TGTAAGTAGAGAAGAGTCTGTATAACCGTGGTGGACTTCCCGCCGAAATCAAACTGCGGGAAGTCAACCACGGTTACATTGGCTCTTCTTACT | ssDNA | Generation of *35S:shc-A12G-NbSu* |
| AC-975 | AATGAGTAAGAAGAGCCAATGTAACCGTGGTTGACTTCCCGCAGTTTGATTTCGGCGGGAAGTCCACCACGGTTATACAGACTCTTCTCTACT | ssDNA | |
| AC-976 | TGTAAGTAGAGAAGAATCAGTATAACCGTGGTGGACTTCCCGCCGAAATCAAACTGCGGGAAGTCAACCACGGTTACATTGGCTCTTCTTACT | ssDNA | Generation of *35S:shc-U15A-NbSu* |

| | | | |
|---|---|---|---|
| AC-977 | AATGAGTAAGAAGAGCCAATGTAACCGTGGTTGACTTCCCGCAGTTTGATTTCGGCGGGAAGTCCACCACGGTTATACTGATTCTTCTCTACT | ssDNA | |
| AC-982 | TGTATAACCGTGGTGGACTTCCCGCCGAAATCAAACGGCGGGAAGTCAACCACGGTTA | ssDNA | Generation of *35S:shc-U51G-NbSu* |
| AC-983 | AATGTAACCGTGGTTGACTTCCCGCCGTTTGATTTCGGCGGGAAGTCCACCACGGTTA | ssDNA | |
| AC-1077 | TGTAAGTAGAGAAGAATCGGTATAACCGTGGTGGACTTCCCGCCGAAATCAAACTGCGGGAAGTCAACCACGGTTACATCGGCTCTTCTTACT | ssDNA | Generation of *35S:shc-U15G/U76C-NbSu* |
| AC-1078 | AATGAGTAAGAAGAGCCGATGTAACCGTGGTTGACTTCCCGCAGTTTGATTTCGGCGGGAAGTCCACCACGGTTATACCGATTCTTCTCTACT | ssDNA | |
| AC-1085 | TGTAAGTAGAGAAGAATCGGTATAAACCGCGGGTTCCTAACAGCGAAATCAAACTCTGTTAGGAAACCGCGGTTTACATCGGCTCTTCTTACT | ssDNA | Generation of *35S:shc-U15G/U76C-NbDXS* |
| AC-1086 | AATGAGTAAGAAGAGCCGATGTAAACCGCGGTTTCCTAACAGAGTTTGATTTCGCTGTTAGGAACCCGCGGTTTATACCGATTCTTCTCTACT | ssDNA | |
| AC-1087 | TGTAAGTAGAGAAGAGTCTGTGTAAACCGCGGGTTCCTAACAGCGAAATCAAACTCTGTTAGGAAACCGCGGTTTACATTGGCTCTTCTTACT | ssDNA | Generation of *35S:shc-A12G/A18G-NbDXS* |
| AC-1088 | AATGAGTAAGAAGAGCCAATGTAAACCGCGGTTTCCTAACAGAGTTTGATTTCGCTGTTAGGAACCCGCGGTTTACACAGACTCTTCTCTACT | ssDNA | |
| AC-1114 | TGTATAACCGTGGTGGACTTCCCGCGGAAATCAAACCGCGGGAAGTCAACCACGGTTA | ssDNA | Generation of *35S:shc-C40G/U51C-NbSu* |
| AC-1115 | AATGTAACCGTGGTTGACTTCCCGCGGTTTGATTTCCGCGGGAAGTCCACCACGGTTA | ssDNA | |
| AC-1116 | TGTATAACCGTGGTGGACTTCCCGCAGAAATCAAACTGCGGGAAGTCAACCACGGTTA | ssDNA | Generation of *35S:shc-C40A-NbSu* |
| AC-1117 | AATGTAACCGTGGTTGACTTCCCGCAGTTTGATTTCTGCGGGAAGTCCACCACGGTTA | ssDNA | |
| AC-1118 | TGTATAACCGTGGTGGACTTCCCGCTGAAATCAAACAGCGGGAAGTCAACCACGGTTA | ssDNA | Generation of *35S:shc-C40U/U51A-NbSu* |
| AC-1119 | AATGTAACCGTGGTTGACTTCCCGCTGTTTGATTTCAGCGGGAAGTCCACCACGGTTA | ssDNA | |
| AC-1120 | TGTATAACCGTGGTGGACTTCCCGCTGAAATCAAACGGCGGGAAGTCAACCACGGTTA | ssDNA | Generation of *35S:shc-C40U/U51G-NbSu* |
| AC-1121 | AATGTAACCGTGGTTGACTTCCCGCCGTTTGATTTCAGCGGGAAGTCCACCACGGTTA | ssDNA | |
| AC-1122 | TGTATAACCGTGGTGGACTTCCCGCGGAAATCAAACTGCGGGAAGTCAACCACGGTTA | ssDNA | Generation of *35S:shc-C40G-NbSu* |
| AC-1123 | AATGTAACCGTGGTTGACTTCCCGCAGTTTGATTTCCGCGGGAAGTCCACCACGGTTA | ssDNA | |
| AC-1124 | TGTATAAACCGCGGGTTCCTAACAGGGAAATCAAACCCTGTTAGGAAACCGCGGTTTA | ssDNA | Generation of *35S:shc-C40G/U51C-NbDXS* |
| AC-1125 | AATGTAAACCGCGGTTTCCTAACAGGGTTTGATTTCCCTGTTAGGAACCCGCGGTTTA | ssDNA | |
| AC-1126 | TGTATAAACCGCGGGTTCCTAACAGAGAAATCAAACTCTGTTAGGAAACCGCGGTTTA | ssDNA | Generation of *35S:shc-C40A-NbDXS* |
| AC-1127 | AATGTAAACCGCGGTTTCCTAACAGAGTTTGATTTCTCTGTTAGGAACCCGCGGTTTA | ssDNA | |
| AC-1128 | TGTATAAACCGCGGGTTCCTAACAGTGAAATCAAACACTGTTAGGAAACCGCGGTTTA | ssDNA | Generation of *35S:shc-C40U/U51A-NbDXS* |
| AC-1129 | AATGTAAACCGCGGTTTCCTAACAGTGTTTGATTTCACTGTTAGGAACCCGCGGTTTA | ssDNA | |
| AC-1130 | TGTATAAACCGCGGGTTCCTAACAGTGAAATCAAACGCTGTTAGGAAACCGCGGTTTA | ssDNA | Generation of *35S:shc-C40U/U51G-NbDXS* |
| AC-1131 | AATGTAAACCGCGGTTTCCTAACAGCGTTTGATTTCACTGTTAGGAACCCGCGGTTTA | ssDNA | |
| AC-1132 | TGTATAAACCGCGGGTTCCTAACAGGGAAATCAAACTCTGTTAGGAAACCGCGGTTTA | ssDNA | Generation of *35S:shc-C40G-NbDXS* |
| AC-1133 | AATGTAAACCGCGGTTTCCTAACAGAGTTTGATTTCCCTGTTAGGAACCCGCGGTTTA | ssDNA | |

| | | | |
|---|---|---|---|
| AC-1134 | TGTAAGTAGAGAAGAATCCGTATAACCGTGGTGGACTTCCCGCCGAAATCAAACTGCGGGAAGTCAACCACGGTTACATGGGCTCTTCTTACT | ssDNA | Generation of *35S:shc-U15C/U76G-NbSu* |
| AC-1135 | AATGAGTAAGAAGAGCCCATGTAACCGTGGTTGACTTCCCGCAGTTTGATTTCGGCGGGAAGTCCACCACGGTTATACGGATTCTTCTCTACT | ssDNA | |
| AC-1136 | TGTAAGTAGAGAAGAATCTGTATAACCGTGGTGGACTTCCCGCCGAAATCAAACTGCGGGAAGTCAACCACGGTTACATAGGCTCTTCTTACT | ssDNA | Generation of *35S:shc-U76A-NbSu* |
| AC-1137 | AATGAGTAAGAAGAGCCTATGTAACCGTGGTTGACTTCCCGCAGTTTGATTTCGGCGGGAAGTCCACCACGGTTATACAGATTCTTCTCTACT | ssDNA | |
| AC-1138 | TGTAAGTAGAGAAGAATCGGTATAACCGTGGTGGACTTCCCGCCGAAATCAAACTGCGGGAAGTCAACCACGGTTACATTGGCTCTTCTTACT | ssDNA | Generation of *35S:shc-U15G-NbSu* |
| AC-1139 | AATGAGTAAGAAGAGCCAATGTAACCGTGGTTGACTTCCCGCAGTTTGATTTCGGCGGGAAGTCCACCACGGTTATACCGATTCTTCTCTACT | ssDNA | |
| AC-1140 | TGTAAGTAGAGAAGAATCTGTATAACCGTGGTGGACTTCCCGCCGAAATCAAACTGCGGGAAGTCAACCACGGTTACATGGGCTCTTCTTACT | ssDNA | Generation of *35S:shc-U76G-NbSu* |
| AC-1141 | AATGAGTAAGAAGAGCCCATGTAACCGTGGTTGACTTCCCGCAGTTTGATTTCGGCGGGAAGTCCACCACGGTTATACAGATTCTTCTCTACT | ssDNA | |
| AC-1142 | TGTAAGTAGAGAAGAATCCGTATAAACCGCGGGTTCCTAACAGCGAAATCAAACTCTGTTAGGAAACCGCGGTTTACATGGGCTCTTCTTACT | ssDNA | Generation of *35S:shc-U15C/U76G-NbDXS* |
| AC-1143 | AATGAGTAAGAAGAGCCCATGTAAACCGCGGTTTCCTAACAGAGTTTGATTTCGCTGTTAGGAACCCGCGGTTTATACGGATTCTTCTCTACT | ssDNA | |
| AC-1144 | TGTAAGTAGAGAAGAATCTGTATAAACCGCGGGTTCCTAACAGCGAAATCAAACTCTGTTAGGAAACCGCGGTTTACATAGGCTCTTCTTACT | ssDNA | Generation of *35S:shc-U76A-NbDXS* |
| AC-1145 | AATGAGTAAGAAGAGCCTATGTAAACCGCGGTTTCCTAACAGAGTTTGATTTCGCTGTTAGGAACCCGCGGTTTATACAGATTCTTCTCTACT | ssDNA | |
| AC-1146 | TGTAAGTAGAGAAGAATCGGTATAAACCGCGGGTTCCTAACAGCGAAATCAAACTCTGTTAGGAAACCGCGGTTTACATTGGCTCTTCTTACT | ssDNA | Generation of *35S:shc-U15G-NbDXS* |
| AC-1147 | AATGAGTAAGAAGAGCCAATGTAAACCGCGGTTTCCTAACAGAGTTTGATTTCGCTGTTAGGAACCCGCGGTTTATACCGATTCTTCTCTACT | ssDNA | |
| AC-1148 | TGTAAGTAGAGAAGAATCTGTATAAACCGCGGGTTCCTAACAGCGAAATCAAACTCTGTTAGGAAACCGCGGTTTACATGGGCTCTTCTTACT | ssDNA | Generation of *35S:shc-U76G-NbDXS* |
| AC-1149 | AATGAGTAAGAAGAGCCCATGTAAACCGCGGTTTCCTAACAGAGTTTGATTTCGCTGTTAGGAACCCGCGGTTTATACAGATTCTTCTCTACT | ssDNA | |
| AC-1150 | TGTAAGTAGAGAAGACTCTGTATAACCGTGGTGGACTTCCCGCCGAAATCAAACTGCGGGAAGTCAACCACGGTTACATTGGGTCTTCTTACT | ssDNA | Generation of *35S:shc-A12C/C79G-NbSu* |
| AC-1151 | AATGAGTAAGAAGACCCAATGTAACCGTGGTTGACTTCCCGCAGTTTGATTTCGGCGGGAAGTCCACCACGGTTATACAGAGTCTTCTCTACT | ssDNA | |
| AC-1152 | TGTAAGTAGAGAAGAATCTGTATAACCGTGGTGGACTTCCCGCCGAAATCAAACTGCGGGAAGTCAACCACGGTTACATTGGTTCTTCTTACT | ssDNA | Generation of *35S:shc-C79U-NbSu* |
| AC-1153 | AATGAGTAAGAAGAACCAATGTAACCGTGGTTGACTTCCCGCAGTTTGATTTCGGCGGGAAGTCCACCACGGTTATACAGATTCTTCTCTACT | ssDNA | |
| AC-1154 | TGTAAGTAGAGAAGATTCTGTATAACCGTGGTGGACTTCCCGCCGAAATCAAACTGCGGGAAGTCAACCACGGTTACATTGGATCTTCTTACT | ssDNA | Generation of *35S:shc-A12U/C79A-NbSu* |
| AC-1155 | AATGAGTAAGAAGATCCAATGTAACCGTGGTTGACTTCCCGCAGTTTGATTTCGGCGGGAAGTCCACCACGGTTATACAGAATCTTCTCTACT | ssDNA | |

| | | | |
|---|---|---|---|
| AC-1156 | TGTAAGTAGAGAAGAGTCTGTATAACCGTGGTGGACTTCCCGCCGAAATCAAACTGCGGGAAGTCAACCACGGTTACATTGGTTCTTCTTACT | ssDNA | Generation of *35S:shc-A12G/C79U-NbSu* |
| AC-1157 | AATGAGTAAGAAGAACCAATGTAACCGTGGTTGACTTCCCGCAGTTTGATTTCGGCGGGAAGTCCACCACGGTTATACAGACTCTTCTCTACT | ssDNA | |
| AC-1158 | TGTAAGTAGAGAAGATTCTGTATAACCGTGGTGGACTTCCCGCCGAAATCAAACTGCGGGAAGTCAACCACGGTTACATTGGGTCTTCTTACT | ssDNA | Generation of *35S:shc-A12U/C79G-NbSu* |
| AC-1159 | AATGAGTAAGAAGACCCAATGTAACCGTGGTTGACTTCCCGCAGTTTGATTTCGGCGGGAAGTCCACCACGGTTATACAGAATCTTCTCTACT | ssDNA | |
| AC-1160 | TGTAAGTAGAGAAGACTCTGTATAAACCGCGGGTTCCTAACAGCGAAATCAAACTCTGTTAGGAAACCGCGGTTTACATTGGGTCTTCTTACT | ssDNA | Generation of *35S:shc-A12C/C79G* |
| AC-1161 | AATGAGTAAGAAGACCCAATGTAAACCGCGGTTTCCTAACAGAGTTTGATTTCGCTGTTAGGAACCCGCGGTTTATACAGAGTCTTCTCTACT | ssDNA | |
| AC-1162 | TGTAAGTAGAGAAGAATCTGTATAAACCGCGGGTTCCTAACAGCGAAATCAAACTCTGTTAGGAAACCGCGGTTTACATTGGTTCTTCTTACT | ssDNA | Generation of *35S:shc-C79U-NbDXS* |
| AC-1163 | AATGAGTAAGAAGAACCAATGTAAACCGCGGTTTCCTAACAGAGTTTGATTTCGCTGTTAGGAACCCGCGGTTTATACAGATTCTTCTCTACT | ssDNA | |
| AC-1164 | TGTAAGTAGAGAAGATTCTGTATAAACCGCGGGTTCCTAACAGCGAAATCAAACTCTGTTAGGAAACCGCGGTTTACATTGGATCTTCTTACT | ssDNA | Generation of *35S:shc-A12U/C79A-NbDXS* |
| AC-1165 | AATGAGTAAGAAGATCCAATGTAAACCGCGGTTTCCTAACAGAGTTTGATTTCGCTGTTAGGAACCCGCGGTTTATACAGAATCTTCTCTACT | ssDNA | |
| AC-1166 | TGTAAGTAGAGAAGAGTCTGTATAAACCGCGGGTTCCTAACAGCGAAATCAAACTCTGTTAGGAAACCGCGGTTTACATTGGTTCTTCTTACT | ssDNA | Generation of *35S:shc-A12G/C79U-NbDXS* |
| AC-1167 | AATGAGTAAGAAGAACCAATGTAAACCGCGGTTTCCTAACAGAGTTTGATTTCGCTGTTAGGAACCCGCGGTTTATACAGACTCTTCTCTACT | ssDNA | |
| AC-1168 | TGTAAGTAGAGAAGATTCTGTATAAACCGCGGGTTCCTAACAGCGAAATCAAACTCTGTTAGGAAACCGCGGTTTACATTGGGTCTTCTTACT | ssDNA | Generation of *35S:shc-A12U/C79G-NbDXS* |
| AC-1169 | AATGAGTAAGAAGACCCAATGTAAACCGCGGTTTCCTAACAGAGTTTGATTTCGCTGTTAGGAACCCGCGGTTTATACAGAATCTTCTCTACT | ssDNA | |
| AC-1180 | TGTATGCGCTTGCTGAGTTTCCCCCCGAAATCAAACTGGGGGAAACTAAGCAAGCGCA | ssDNA | Generation of *35S:shc-* $GUS_{At}$ |
| AC-1181 | AATGTGCGCTTGCTTAGTTTCCCCCAGTTTGATTTCGGGGGGAAACTCAGCAAGCGCA | ssDNA | |
| AC-1237 | TGTAAGTAGAGAAGAGTCGGTATAAACCGCGGGTTCCTAACAGCGAAATCAAACTCTGTTAGGAAACCGCGGTTTACATCGGCTCTTCTTACT | ssDNA | Generation of *35S:shc-A12G/U15G-NbDXS* |
| AC-1238 | AATGAGTAAGAAGAGCCGATGTAAACCGCGGTTTCCTAACAGAGTTTGATTTCGCTGTTAGGAACCCGCGGTTTATACCGACTCTTCTCTACT | ssDNA | |
| AC-1239 | TGTAAGTAGAGAAGAATCGGTGTAAACCGCGGGTTCCTAACAGCGAAATCAAACTCTGTTAGGAAACCGCGGTTTACATCGGCTCTTCTTACT | ssDNA | Generation of *35S:shc-U15G/A18G-NbDXS* |
| AC-1240 | AATGAGTAAGAAGAGCCGATGTAAACCGCGGTTTCCTAACAGAGTTTGATTTCGCTGTTAGGAACCCGCGGTTTACACCGATTCTTCTCTACT | ssDNA | |
| AC-1241 | TGTAAGTAGAGAAGAGTCGGTGTAAACCGCGGGTTCCTAACAGCGAAATCAAACTCTGTTAGGAAACCGCGGTTTACATCGGCTCTTCTTACT | ssDNA | Generation of *35S:shc-A12G/U15G/A18G-NbDXS* |
| AC-1242 | AATGAGTAAGAAGAGCCGATGTAAACCGCGGTTTCCTAACAGAGTTTGATTTCGCTGTTAGGAACCCGCGGTTTACACCGACTCTTCTCTACT | ssDNA | |
| AC-1268 | CACCAGTAGAGAAGAATCTGTGAGAGACCggtctcAcattggctcttcttact | ssDNA | Generation of *pENTR-BS-AtMIR390a-A18G-* |

| | | | |
|---|---|---|---|
| AC-1269 | agtaagaagagccaatgTgagaccGGTCTCTCACAGATTCTTCTCTAC TGGTG | ssDNA | BB and pMDC32B-*BS-AtMIR390a-A18G-BB* |
| AC-1270 | CACCTGTGAGAGACCATTAGGCACCCC | ssDNA | Generation of *pENTR-BS-AtMIR390a-A18G-B/c* and *pMDC32B-BS-AtMIR390a-A18G-B/c* |
| AC-1271 | aatgTgagaccGTCGAGGTGC | ssDNA | |
| AC-1272 | TGTAAAGCTCAGGAGGGATAGCGCCATGATGATCACATT CGTTATCTATTTTTTGGCGCTATCCATCCTGAGTTT | ssDNA | Generation of *35S:AtMIR390a* |
| AC-1273 | AATGAAACTCAGGATGGATAGCGCCAAAAAATAGATAA CGAATGTGATCATCATGGCGCTATCCCTCCTGAGCTT | ssDNA | |
| AC-1274 | TGTAAGTAGAGAAGAATCTGTGAAGCTCAGGAGGGATA GCGCCATGATGATCACATTCGTTATCTATTTTTTGGCGCT ATCCATCCTGAGTTTCATTGGCTCTTCTTACT | ssDNA | Generation of *35S:AtMIR390a-A18G* |
| AC-1275 | AATGAGTAAGAAGAGCCAATGAAACTCAGGATGGATAG CGCCAAAAAATAGATAACGAATGTGATCATCATGGCGC TATCCCTCCTGAGCTTCACAGATTCTTCTCTACT | ssDNA | |
| AC-1276 | TGTGTTGGTTATAAAGGAAGAGGCCCGAAATCAAACTG GCCTCTTCCGTTATAACCAA | ssDNA | Generation of *35S:shc-A18G-AtFT* |
| AC-1277 | AATGTTGGTTATAACGGAAGAGGCCAGTTTGATTTCGGG CCTCTTCCTTTATAACCAA | ssDNA | |
| AC-1278 | TGTGTTAAGTGTCACGGAAATCCCTCGAAATCAAACTAG GGATTTCCTTGACACTTAA | ssDNA | Generation of *35S:shc-A18G-AtCH42* |
| AC-1279 | AATGTTAAGTGTCAAGGAAATCCCTAGTTTGATTTCGAG GGATTTCCGTGACACTTAA | ssDNA | |
| AC-1280 | TGTATTCGCCTTGACCTGATCCCTTCGAAATCAAACTAA GGGATCAGTTCAAGGCGAA | ssDNA | Generation of *35S:shc-AtELF3* |
| AC-1281 | AATGTTCGCCTTGAACTGATCCCTTAGTTTGATTTCGAAG GGATCAGGTCAAGGCGAA | ssDNA | |
| AC-1282 | TGTGTTCGCCTTGACCTGATCCCTTCGAAATCAAACTAA GGGATCAGTTCAAGGCGAA | ssDNA | Generation of *35S:shc-A18G-AtELF3* |
| AC-1286 | TGTAAGTAGAGAAGAATCTGTGTAACCGTGGTGGACTTC CCGCATGATGATCACATTCGTTATCTATTTTTGCGGGAA GTCAACCACGGTTACATTGGCTCTTCTTACT | ssDNA | Generation of *35S:AtMIR390a-A18G-NbSu* |
| AC-1287 | AATGAGTAAGAAGAGCCAATGTAACCGTGGTTGACTTCC CGCAAAAAATAGATAACGAATGTGATCATCATGCGGGA AGTCCACCACGGTTACACAGATTCTTCTCTACT | ssDNA | |
| AC-1288 | TGTAAGTAGAGAAGAATCTGTGTAAACCGCGGGTTCCTA ACAGATGATGATCACATTCGTTATCTATTTTTTCTGTTAG GAAACCGCGGTTTACATTGGCTCTTCTTACT | ssDNA | Generation of *35S:AtMIR390a-A18G-NbDXS* |
| AC-1289 | AATGAGTAAGAAGAGCCAATGTAAACCGCGGTTTCCTA ACAGAAAAAATAGATAACGAATGTGATCATCATCTGTTA GGAACCCGCGGTTTACACAGATTCTTCTCTACT | ssDNA | |

*ssDNA: single-stranded DNA; dsDNA: double-stranded DNA; LNA: locked nucleic acid.

### Text S1

Protocol to design and clone amiRNAs downstream the BS region in *BS-AtMIR390a-A18G-Bsa*I/*ccd*B-based ('B/c') vectors.

#### 1. Selection of the amiRNA sequence

Use the amiRNA Designer tool from the P-SAMS script at https://github.com/carringtonlab/p-sams.

#### 2. Design of amiRNA oligonucleotides

Use amiRNA Designer tool from the P-SAMS script at https://github.com/carringtonlab/p-sams.

2.2.1 Sequence of the *BS-AtMIR390a-A18G* cassette containing the amiRNA

The following FASTA sequence includes amiRNA/amiRNA* sequences inserted in the *BS-AtMIR390a-A18G* precursor sequence downstream the BS region:

**>amiRNA in *BS-AtMIR390a-A18G***

**AGTAGAGAAGAATCTGTGG**$X_1X_2X_3X_4X_5X_6X_7X_8X_9X_{10}X_{11}X_{12}X_{13}X_{14}X_{15}X_{16}X_{17}X_{18}X_{19}X_{20}X_{21}$CGAAATCAAACT$X_1X_2X_1X_2X_3X_4X_5X_6X_7X_8X_9X_{10}X_{11}X_{12}X_{13}X_{14}X_{15}X_{16}X_{17}X_{18}X_{19}$CA**TTGGCTCTTCTTACT**

Where:

-X is a DNA base of the amiRNA sequence, and the subscript number is the base position in the amiRNA 21-mer

-X is a DNA base of the amiRNA* sequence, and the subscript number is the base position in the amiRNA* 21-mer

-**X** is a DNA base of the BS region of the *AtMIR390a* precursor

-X is a DNA base of the *OsMIR390* precursor included in the oligonucleotides required to clone the amiRNA insert in B/c vectors

-G is A18G modification

-X is a DNA base of the *AtMIR390a* precursor included in the oligonucleotides required to clone the amiRNA insert in B/c vectors

-X is a DNA base of the *OsMIR390a* precursor that may be modified to preserve the authentic *AtMIR390a* duplex structure

In the sequence above:

-Insert the amiRNA sequence where you see

$X_1X_2X_3X_4X_5X_6X_7X_8X_9X_{10}X_{11}X_{12}X_{13}X_{14}X_{15}X_{16}X_{17}X_{18}X_{19}X_{20}X_{21}$

-Insert the amiRNA* sequence that has to verify the following base-pairing:

```
 X1 X2 X3 X4 X5 X6 X7 X8 X9 X10X11X12X13X14X15X16X17X18 X19 X20X21
 |  |  |  |  |  |  |  |  |  |      |  |  |  |  |  |  |   |   |   |
X19X18X17X16X15X14X13X12X11X10X9 X8 X7 X6 X5 X4 X3 X2 X1  X2 X1
```

Note that:

-In general, **$X_1$=T** for amiRNA association with AGO1. In this case, **$X_{19}$=A**

-Bases $X_{11}$ and $X_9$ DO NOT base-pair to preserve the central bulge of the authentic *AtMIR390a* duplex. The following base-pair rule applies:

-If $X_{11}$=G, then $X_9$=A

-If $X_{11}$=C, then $X_9$=T

-If $X_{11}$=A, then $X_9$=G

-If $X_{11}$=U, then $X_9$=C

2.2.2. Sequence of the amiRNA oligonucleotides

The sequences of the two amiRNA oligonucleotides are:

-Forward oligonucleotide (58 b),

**TGTG**$X_1X_2X_3X_4X_5X_6X_7X_8X_9X_{10}X_{11}X_{12}X_{13}X_{14}X_{15}X_{16}X_{17}X_{18}X_{19}X_{20}X_{21}$CGAAATCAAACT$X_1X_2X_1X_2X_3X_4X_5X_6X_7X_8X_9X_{10}X_{11}X_{12}X_{13}X_{14}X_{15}X_{16}X_{17}X_{18}X_{19}$

-Reverse oligonucleotide (58 b),

**AA**TG$Y_{19}Y_{18}Y_{17}Y_{16}Y_{15}Y_{14}Y_{13}Y_{12}Y_{11}Y_{10}Y_9Y_8Y_7Y_6Y_5Y_4Y_3Y_2Y_1Y_2Y_1$AGTTTGATTTCG$Y_{21}Y_{20}Y_{19}Y_{18}Y_{17}Y_{16}Y_{15}Y_{14}Y_{13}Y_{12}Y_{11}Y_{10}Y_9Y_8Y_7Y_6Y_5Y_4Y_3Y_2Y_1$

Where:

-$X_1X_2X_3X_4X_5X_6X_7X_8X_9X_{10}X_{11}X_{12}X_{13}X_{14}X_{15}X_{16}X_{17}X_{18}X_{19}X_{20}X_{21}$=amiRNA sequence

-$X_1X_2X_3X_4X_5X_6X_7X_8X_9X_{10}X_{11}X_{12}X_{13}X_{14}X_{15}X_{16}X_{17}X_{18}X_{19}$=partial amiRNA* sequence

-$Y_{21}Y_{20}Y_{19}Y_{18}Y_{17}Y_{16}Y_{15}Y_{14}Y_{13}Y_{12}Y_{11}Y_{10}Y_9Y_8Y_7Y_6Y_5Y_4Y_3Y_2Y_1$=amiRNA reverse-complement sequence

-TG$Y_{19}Y_{18}Y_{17}Y_{16}Y_{15}Y_{14}Y_{13}Y_{12}Y_{11}Y_{10}Y_9Y_8Y_7Y_6Y_5Y_4Y_3Y_2Y_1$=amiRNA* reverse-complement sequence

-$X_1X_2$ = *OsMIR390* sequence that may be modified to preserve authentic *OsMIR390a* duplex structure.

-$Y_2Y_1$ = reverse-complement of $X_1X_2$

**Example:**

The sequences of the two oligonucleotides to clone the amiRNA 'amiR-NbSu' (TCCCATTCGATACTGCTCGCC) are:

-Sense oligonucleotide (58 b),

**TGTG**TAACCGTGGTGGACTTCCCGCCGAAATCAAACTGCGGGAAGTCAACCACGGTTA

-Antisense oligonucleotide (58 b),

**AA**TGTAACCGTGGTTGACTTCCCGCAGTTTGATTTCGGCGGGAAGTCCACCACGGTTA

**Note:** *the 58 b long oligonucleotides can be ordered desalted, no purification is required.*

#### 3. Cloning of amiRNA sequence(s) in *BS-AtMIR390a-A18G-B/c*-based vectors

*Notes:*

*-Available BS-AtMIR390a-A18G-B/c vectors are listed in Table I at the end of the section.*

*-BS-AtMIR390a-A18G-B/c-based vectors must be propagated in a* ccd*B resistant* E. coli *strain such as DB3.1.*

-Alternatively, Bsa*I digestion of the B/c vector and subsequent ligation of the amiRNA oligonucleotide insert can be done in separate reactions*

3.1. Oligonucleotide annealing

-Dilute sense oligonucleotide and antisense oligonucleotide in sterile $H_2O$ to a final concentration of 100 µM.

-Prepare Oligo Annealing Buffer:

60 mM Tris-HCl (pH 7.5)

500 mM NaCl

60 mM $MgCl_2$

10 mM DTT

***Note:*** *Prepare 1 ml aliquots of Oligo Annealing Buffer and store at -20$^0$C.*

-Assemble the annealing reaction in a PCR tube as described below:

| | |
|---|---|
| Forward oligonucleotide (100 µM) | 2 µL |
| Reverse oligonucleotide (100 µM) | 2 µL |
| Oligo Annealing Buffer | 46 µL |

| | |
|---|---|
| Total volume | 50 µL |

The final concentration of each oligonucleotide is 4 µM.

-Use a thermocycler to heat the annealing reaction 5 min at 94°C and then cool down (0.05ºC/sec) to 20°C.

-Dilute the annealed oligonucleotides just prior to assembling the digestion-ligation reaction as described below:

| | |
|---|---|
| Annealed oligonucleotides | 3 µL |
| $dH_2O$ | 37 µL |
| Total volume | 40 µL |

The final concentration of each oligonucleotide is 0.15 µM.

***Note**: Do not store the diluted oligonucleotides.*

3.2. Digestion-ligation reaction

- Assemble the digestion-ligation reaction as described below:

| | |
|---|---|
| B/c vector (x ug/uL) | Y µL (50 ng) |
| Diluted annealed oligonucleotides | 1 µL |
| 10x T4 DNA ligase buffer | 1 µL |
| T4 DNA ligase (400 U/µL) | 1 µL |
| *Bsa*I (10U/ µL, NEB) | 1 µL |
| $dH_2O$ | to 10 µL |
| Total volume | 10 µL |

Prepare a negative control reaction lacking *Bsa*I.

-Mix the reactions by pipetting. Incubate the reactions at room temperature for 5 minutes at 37°C.

3.3. *E. coli* transformation and analysis of transformants

-Transform 1-5 ul of the digestion-ligation reaction into an *E. coli* strain that doesn't have *ccd*B resistance (e.g. DH10B, TOP10, …) to do counter-selection.

-Pick two colonies/construct, grow LB-Kan (100 mg/ml) cultures and purify plasmids.

-Sequence with appropriate primers: M13-F (CCCAGTCACGACGTTGTAAAACGACGG) and M13-R (CAGAGCTGCCAGGAAACAGCTATGACC) for *pENTR*-based vectors; attB1 (ACAAGTTTGTACAAAAAAGCAGGCT) and attB2 (ACCACTTTGTACAAGAAAGCTGGGT) primers for *pMDC32B*-based vectors).

**Table I**: *Bsa*I/*ccd*B-based ('B/c') vectors for direct cloning of amiRNAs downstream the BS region in *AtMIR390a-A18G* precursor.

| Vector | Small RNA expressed | Bacterial antibiotic resistance | Plant antibiotic resistance | GATEWAY use | Backbone | Promoter of syn-tasiRNA cassette | Terminator of syn-tasiRNA cassette | Plant species tested |
|---|---|---|---|---|---|---|---|---|
| *pENTR-BS-AtMIR390a-A18G-B/c* | – | Kanamycin | – | Donor | *pENTR* | – | – | – |
| *pMDC32B-BS-AtMIR390a-A18GB/c* | amiRNA | Kanamycin Hygromycin | Hygromycin | – | *pMDC32* | *CaMV* 2x35S | *Nos* | *A. thaliana* *N. benthamiana* |

**Text S2.**

FASTA sequences of miRNA/amiRNA-producing precursors.

AtMIR390a
OsMIR390
miRNA/amiRNA
miRNA*/amiRNA*
N and N: mutations tested
N is a DNA base of the *OsMIR390a* precursor that may be modified to preserve the authentic *AtMIR390a* duplex structure

>AtMIR390a
AGTAGAGAAGAATCTGTAAAGCTCAGGAGGGATAGCGCCATGATGATCACATTCGTTATCTATTTTTTGGCGCTA
TCCATCCTGAGTTTCATTGGCTCTTCTTACT

>AtMIR390a-A18G
AGTAGAGAAGAATCTGTGAAGCTCAGGAGGGATAGCGCCATGATGATCACATTCGTTATCTATTTTTTGGCGCTA
TCCATCCTGAGTTTCATTGGCTCTTCTTACT

>AtMIR390a-GUS$_{Nb}$
AGTAGAGAAGAATCTGTATCTTGTAACGCGCTTTCCCAGATGATGATCACATTCGTTATCTATTTTTTCTGGGAA
AGCTCGTTACAAGACATTGGCTCTTCTTACT

>AtMIR390a-NbSu
AGTAGAGAAGAATCTGTATAACCGTGGTGGACTTCCCGCATGATGATCACATTCGTTATCTATTTTTTGCGGGAA
GTCAACCACGGTTACATTGGCTCTTCTTACT

>AtMIR390a-A18G-NbSu
AGTAGAGAAGAATCTGTGTAACCGTGGTGGACTTCCCGCATGATGATCACATTCGTTATCTATTTTTTGCGGGAA
GTCAACCACGGTTACATTGGCTCTTCTTACT

>AtMIR390a-NbDXS
AGTAGAGAAGAATCTGTATAAACCGCGGGTTCCTAACAGATGATGATCACATTCGTTATCTATTTTTTCTGTTAG
GAAACCGCGGTTTACATTGGCTCTTCTTACT

>AtMIR390a-A18G-NbDXS
AGTAGAGAAGAATCTGTGTAAACCGCGGGTTCCTAACAGATGATGATCACATTCGTTATCTATTTTTTCTGTTAG
GAAACCGCGGTTTACATTGGCTCTTCTTACT

>shc-GUS$_{Nb}$
AGTAGAGAAGAATCTGTATCTTGTAACGCGCTTTCCCAGCGAAATCAAACTCTGGGAAAGCTCGTTACAAGACAT
TGGCTCTTCTTACT

>shc-NbSu
AGTAGAGAAGAATCTGTATAACCGTGGTGGACTTCCCGCCGAAATCAAACTGCGGGAAGTCAACCACGGTTACAT
TGGCTCTTCTTACT

>shc-A18G-NbSu
AGTAGAGAAGAATCTGTGTAACCGTGGTGGACTTCCCGCCGAAATCAAACTGCGGGAAGTCAACCACGGTTACAT
TGGCTCTTCTTACT

>shc-A18U/C73A-NbSu
AGTAGAGAAGAATCTGTTTAACCGTGGTGGACTTCCCGCCGAAATCAAACTGCGGGAAGTCAACCACGGTTAAAT
TGGCTCTTCTTACT

>shc-A18U/C73G-NbSu

AGTAGAGAAGAATCTGTTTAACCGTGGTGGACTTCCCGCCGAAATCAAACTGCGGGAAGTCAACCACGGTTAGAT
TGGCTCTTCTTACT

>shc-A18G/C73U-NbSu
AGTAGAGAAGAATCTGTGTAACCGTGGTGGACTTCCCGCCGAAATCAAACTGCGGGAAGTCAACCACGGTTATAT
TGGCTCTTCTTACT

>shc-A18C/C73G-NbSu
AGTAGAGAAGAATCTGTCTAACCGTGGTGGACTTCCCGCCGAAATCAAACTGCGGGAAGTCAACCACGGTTAGAT
TGGCTCTTCTTACT

>shc-NbDXS
AGTAGAGAAGAATCTGTATAAACCGCGGGTTCCTAACAGCGAAATCAAACTCTGTTAGGAAACCGCGGTTTACAT
TGGCTCTTCTTACT

>shc-A18G-NbDXS
AGTAGAGAAGAATCTGTGTAAACCGCGGGTTCCTAACAGCGAAATCAAACTCTGTTAGGAAACCGCGGTTTACAT
TGGCTCTTCTTACT

>shc-C73U-NbDXS
AGTAGAGAAGAATCTGTATAAACCGCGGGTTCCTAACAGCGAAATCAAACTCTGTTAGGAAACCGCGGTTTATAT
TGGCTCTTCTTACT

>shc-A18U/C73A-NbDXS
AGTAGAGAAGAATCTGTTTAAACCGCGGGTTCCTAACAGCGAAATCAAACTCTGTTAGGAAACCGCGGTTTAAAT
TGGCTCTTCTTACT

>shc-A18U/C73G-NbDXS
AGTAGAGAAGAATCTGTTTAAACCGCGGGTTCCTAACAGCGAAATCAAACTCTGTTAGGAAACCGCGGTTTAGAT
TGGCTCTTCTTACT

>shc-A18G/C73U-NbDXS
AGTAGAGAAGAATCTGTGTAAACCGCGGGTTCCTAACAGCGAAATCAAACTCTGTTAGGAAACCGCGGTTTATAT
TGGCTCTTCTTACT

>shc-A18C/C73G-NbDXS
AGTAGAGAAGAATCTGTCTAAACCGCGGGTTCCTAACAGCGAAATCAAACTCTGTTAGGAAACCGCGGTTTAGAT
TGGCTCTTCTTACT

>shc-C40G/U51C-NbSu
AGTAGAGAAGAATCTGTATAACCGTGGTGGACTTCCCGCGGAAATCAAACCGCGGGAAGTCAACCACGGTTACAT
TGGCTCTTCTTACT

>shc-U51G-NbSu
AGTAGAGAAGAATCTGTATAACCGTGGTGGACTTCCCGCCGAAATCAAACGGCGGGAAGTCAACCACGGTTACAT
TGGCTCTTCTTACT

>shc-C40A-NbSu
AGTAGAGAAGAATCTGTATAACCGTGGTGGACTTCCCGCAGAAATCAAACTGCGGGAAGTCAACCACGGTTACAT
TGGCTCTTCTTACT

>shc-C40U/U51A-NbSu
AGTAGAGAAGAATCTGTATAACCGTGGTGGACTTCCCGCTGAAATCAAACAGCGGGAAGTCAACCACGGTTACAT
TGGCTCTTCTTACT

>shc-C40U/U51G-NbSu
AGTAGAGAAGAATCTGTATAACCGTGGTGGACTTCCCGCTGAAATCAAACGGCGGGAAGTCAACCACGGTTACAT
TGGCTCTTCTTACT

>shc-C40G-NbSu
AGTAGAGAAGAATCTGTATAACCGTGGTGGACTTCCCGC**G**GAAATCAAACTGCGGGAAGTCAACCACGGTTACAT
TGGCTCTTCTTACT

>shc-C40G/U51C-NbDXS
AGTAGAGAAGAATCTGTATAAACCGCGGGTTCCTAACAG**G**GAAATCAAAC**C**CTGTTAGGAAACCGCGGTTTACAT
TGGCTCTTCTTACT

>shc-U51G-NbDXS
AGTAGAGAAGAATCTGTATAAACCGCGGGTTCCTAACAGCGAAATCAAAC**G**CTGTTAGGAAACCGCGGTTTACAT
TGGCTCTTCTTACT

>shc-C40A-NbDXS
AGTAGAGAAGAATCTGTATAAACCGCGGGTTCCTAACAG**A**GAAATCAAACTCTGTTAGGAAACCGCGGTTTACAT
TGGCTCTTCTTACT

>shc-C40U/U51A-NbDXS
AGTAGAGAAGAATCTGTATAAACCGCGGGTTCCTAACAGTGAAATCAAAC**A**CTGTTAGGAAACCGCGGTTTACAT
TGGCTCTTCTTACT

>shc-C40U/U51G-NbDXS
AGTAGAGAAGAATCTGTATAAACCGCGGGTTCCTAACAGTGAAATCAAAC**G**CTGTTAGGAAACCGCGGTTTACAT
TGGCTCTTCTTACT

>shc-C40G-NbDXS
AGTAGAGAAGAATCTGTATAAACCGCGGGTTCCTAACAG**G**GAAATCAAACTCTGTTAGGAAACCGCGGTTTACAT
TGGCTCTTCTTACT

>shc-A12G-NbSu
AGTAGAGAAGA**G**TCTGTATAACCGTGGTGGACTTCCCGCCGAAATCAAACTGCGGGAAGTCAACCACGGTTACAT
TGGCTCTTCTTACT

>shc-C79U-NbSu
AGTAGAGAAGAATCTGTATAACCGTGGTGGACTTCCCGCCGAAATCAAACTGCGGGAAGTCAACCACGGTTACAT
TGG**T**TCTTCTTACT

>shc-A12U/C79A-NbSu
AGTAGAGAAGA**T**TCTGTATAACCGTGGTGGACTTCCCGCCGAAATCAAACTGCGGGAAGTCAACCACGGTTACAT
TGG**A**TCTTCTTACT

>shc-A12U/C79G-NbSu
AGTAGAGAAGA**T**TCTGTATAACCGTGGTGGACTTCCCGCCGAAATCAAACTGCGGGAAGTCAACCACGGTTACAT
TGG**G**TCTTCTTACT

>shc-A12C/C79G-NbSu
AGTAGAGAAGA**C**TCTGTATAACCGTGGTGGACTTCCCGCCGAAATCAAACTGCGGGAAGTCAACCACGGTTACAT
TGG**G**TCTTCTTACT

>shc-A12G-C79U-NbSu
AGTAGAGAAGA**G**TCTGTATAACCGTGGTGGACTTCCCGCCGAAATCAAACTGCGGGAAGTCAACCACGGTTACAT
TGG**T**TCTTCTTACT

>shc-A12G-NbDXS
AGTAGAGAAGA**G**TCTGTATAAACCGCGGGTTCCTAACAGCGAAATCAAACTCTGTTAGGAAACCGCGGTTTACAT
TGGCTCTTCTTACT

>shc-C79U-NbDXS

AGTAGAGAAGAATCTGTATAAACCGCGGGTTCCTAACAGCGAAATCAAACTCTGTTAGGAAACCGCGGTTTACAT
TGGTTCTTCTTACT

>shc-A12U/C79A-NbDXS
AGTAGAGAAGATTCTGTATAAACCGCGGGTTCCTAACAGCGAAATCAAACTCTGTTAGGAAACCGCGGTTTACAT
TGGATCTTCTTACT

>shc-A12U/C79G-NbDXS
AGTAGAGAAGATTCTGTATAAACCGCGGGTTCCTAACAGCGAAATCAAACTCTGTTAGGAAACCGCGGTTTACAT
TGGGTCTTCTTACT

>shc-A12C/C79G-NbDXS
AGTAGAGAAGACTCTGTATAAACCGCGGGTTCCTAACAGCGAAATCAAACTCTGTTAGGAAACCGCGGTTTACAT
TGGGTCTTCTTACT

>shc-A12G-C79U-NbDXS
AGTAGAGAAGAGTCTGTATAAACCGCGGGTTCCTAACAGCGAAATCAAACTCTGTTAGGAAACCGCGGTTTACAT
TGGTTCTTCTTACT

>shc-U15G/U76C-NbSu
AGTAGAGAAGAATCGGTATAACCGTGGTGGACTTCCCGCCGAAATCAAACTGCGGGAAGTCAACCACGGTTACAT
CGGCTCTTCTTACT

>shc-U15C/U76G-NbSu
AGTAGAGAAGAATCCGTATAACCGTGGTGGACTTCCCGCCGAAATCAAACTGCGGGAAGTCAACCACGGTTACAT
GGGCTCTTCTTACT

>shc-U15A-NbSu
AGTAGAGAAGAATCAGTATAACCGTGGTGGACTTCCCGCCGAAATCAAACTGCGGGAAGTCAACCACGGTTACAT
TGGCTCTTCTTACT

>shc-U76A-NbSu
AGTAGAGAAGAATCTGTATAACCGTGGTGGACTTCCCGCCGAAATCAAACTGCGGGAAGTCAACCACGGTTACAT
AGGCTCTTCTTACT

>shc-U76G-NbSu
AGTAGAGAAGAATCTGTATAACCGTGGTGGACTTCCCGCCGAAATCAAACTGCGGGAAGTCAACCACGGTTACAT
GGGCTCTTCTTACT

>shc-U15G-NbSu
AGTAGAGAAGAATCTGTATAACCGTGGTGGACTTCCCGCCGAAATCAAACGGCGGGAAGTCAACCACGGTTACAT
TGGCTCTTCTTACT

>shc-U15G/U76C-NbDXS
AGTAGAGAAGAATCGGTATAAACCGCGGGTTCCTAACAGCGAAATCAAACTCTGTTAGGAAACCGCGGTTTACAT
CGGCTCTTCTTACT

>shc-U15C/U76G-NbDXS
AGTAGAGAAGAATCCGTATAAACCGCGGGTTCCTAACAGCGAAATCAAACTCTGTTAGGAAACCGCGGTTTACAT
GGGCTCTTCTTACT

>shc-U15A-NbDXS
AGTAGAGAAGAATCAGTATAAACCGCGGGTTCCTAACAGCGAAATCAAACTCTGTTAGGAAACCGCGGTTTACAT
TGGCTCTTCTTACT

>shc-U76A-NbDXS
AGTAGAGAAGAATCTGTATAAACCGCGGGTTCCTAACAGCGAAATCAAACTCTGTTAGGAAACCGCGGTTTACAT
AGGCTCTTCTTACT

>shc-U76G-NbDXS
AGTAGAGAAGAATCTGTATAAACCGCGGGTTCCTAACAGCGAAATCAAACTCTGTTAGGAAACCGCGGTTTACAT
**G**GGCTCTTCTTACT

>shc-U15G-NbDXS
AGTAGAGAAGAATC**G**GTATAAACCGCGGGTTCCTAACAGCGAAATCAAACTCTGTTAGGAAACCGCGGTTTACAT
TGGCTCTTCTTACT

>shc-A12G-U15G-NbDXS
AGTAGAGAAGA**G**TC**G**GTATAAACCGCGGGTTCCTAACAGCGAAATCAAACTCTGTTAGGAAACCGCGGTTTACAT
**C**GGCTCTTCTTACT

>shc-A12G/A18G-NbDXS
AGTAGAGAAGA**G**TCTGT**G**TAAACCGCGGGTTCCTAACAGCGAAATCAAACTCTGTTAGGAAACCGCGGTTTACAT
TGGCTCTTCTTACT

>shc-U15G/A18G-NbDXS
AGTAGAGAAGAATC**G**GT**G**TAAACCGCGGGTTCCTAACAGCGAAATCAAACTCTGTTAGGAAACCGCGGTTTACAT
**C**GGCTCTTCTTACT

>shc-A12G/U15G/A18G-NbDXS
AGTAGAGAAGA**G**TC**G**GT**G**TAAACCGCGGGTTCCTAACAGCGAAATCAAACTCTGTTAGGAAACCGCGGTTTACAT
**C**GGCTCTTCTTACT

>shc-$GUS_{At}$
AGTAGAGAAGAATCTGTATGCGCTTGCTGAGTTTCCCCCGAAATCAAACTGGGGGAAACTAAGCAAGCGCACAT
TGGCTCTTCTTACT

>shc-AtCH42
AGTAGAGAAGAATCTGTATTAAGTGTCACGGAAATCCCTCGAAATCAAACTAGGGATTTCCTTGACACTTAACAT
TGGCTCTTCTTACT

>shc-A18G-AtCH42
AGTAGAGAAGAATCTGT**G**TTAAGTGTCACGGAAATCCCTCGAAATCAAACTAGGGATTTCCTTGACACTTAACAT
TGGCTCTTCTTACT

>shc-AtFT
AGTAGAGAAGAATCTGTATTGGTTATAAAGGAAGAGGCCCGAAATCAAACTGGCCTCTTCCGTTATAACCAACAT
TGGCTCTTCTTACT

>shc-A18G-AtFT
AGTAGAGAAGAATCTGT**G**TTCGCCTTGACCTGATCCCTTCGAAATCAAACTAAGGGATCAGTTCAAGGCGAACAT
TGGCTCTTCTTACT

>shc-AtELF3
AGTAGAGAAGAATCTGTATTCGCCTTGACCTGATCCCTTCGAAATCAAACTAAGGGATCAGTTCAAGGCGAACAT
TGGCTCTTCTTACT

>shc-A18G-AtELF3
AGTAGAGAAGAATCTGT**G**TTCGCCTTGACCTGATCCCTTCGAAATCAAACTAAGGGATCAGTTCAAGGCGAACAT
TGGCTCTTCTTACT

#### Text S3.

DNA sequence of *Bsa*I-*ccd*B-based (B/c) vectors used for direct cloning of amiRNAs in *MIR390*-based *shc* precursors.

**>*pENTR-BS-AtMIR390a-A18G-B/c* (4076 bp)**
CTTTCCTGCGTTATCCCCTGATTCTGTGGATAACCGTATTACCGCCTTTGAGTGAGCTGATACCGCTCGCCGCAG
CCGAACGACCGAGCGCAGCGAGTCAGTGAGCGAGGAAGCGGAAGAGCGCCCAATACGCAAACCGCCTCTCCCCGC
GCGTTGGCCGATTCATTAATGCAGCTGGCACGACAGGTTTCCCGACTGGAAAGCGGGCAGTGAGCGCAACGCAAT
TAATACGCGTACCGCTAGCCAGGAAGAGTTTGTAGAAACGCAAAAAGGCCATCCGTCAGGATGGCCTTCTGCTTA
GTTTGATGCCTGGCAGTTTATGGCGGGCGTCCTGCCCGCCACCCTCCGGGCCGTTGCTTCACAACGTTCAAATCC
GCTCCCGGCGGATTTGTCCTACTCAGGAGAGCGTTCACCGACAAACAACAGATAAAACGAAAGGCCCAGTCTTCC
GACTGAGCCTTTCGTTTTATTTGATGCCTGGCAGTTCCCTACTCTCGCGTTAACGCTAGCATGGATGTTTTCCCA
GTCACGACGTTGTAAAACGACGGCCAGTCTTAAGCTCGGGCCCCAAATAATGATTTTATTTTGACTGATAGTGAC
CTGTTCGTTGCAACAAATTGATGAGCAATGCTTTTTTATAATGCCAACTTTGTACAAAAAAGCAGGCTCCGCGGC
CGCCCCCTTCACCGTAGAGAAGAATCTGT**G**AGAGACCATTAGGCACCCCAGGCTTTACACTTTATGCTTCCGGCT
CGTATAATGTGTGGATTTTGAGTTAGGAGCCGTCGAGATTTTCAGGAGCTAAGGAAGCTAAAatggagaaaaaaa
tcactggatataccaccgttgatatatcccaatggcatcgtaaagaacattttgaggcatttcagtcagttgctc
aatgtacctataaccagaccgttcagctggatattacggcctttttaaagaccgtaaagaaaaataagcacaagt
tttatccggcctttattcacattcttgcccgcctgatgaatgctcatccggagttccgtatggcaatgaaagacg
gtgagctggtgatatgggatagtgttcacccttgttacaccgttttccatgagcaaactgaaacgttttcatcgc
tctggagtgaataccacgacgatttccggcagtttctacacatatattcgcaagatgtggcgtgttacggtgaaa
acctggcctatttccctaaagggtttattgagaatatgtttttcgtctcagccaatccctgggtgagtttcacca
gttttgatttaaacgtggccaatatggacaacttcttcgcccccgttttcaccatgggcaaatattatacgcaag
gcgacaaggtgctgatgccgctggcgattcaggttcatcatgccgtttgtgatggcttccatgtcggcagaatgc
ttaatgaattacaacagtactgcgatgagtggcagggcggggcgtaaACGCGTGGAGCCGGCTTACTAAAAGCCA
GATAACAGTATGCGTATTTGCGCGCTGATTTTTGCGGTATAAGAATATATACTGATATGTATACCCGAAGTATGT
CAAAAAGAGGTATGCTATGAAGCAGCGTATTACAGTGACAGTTGACAGCGACAGCTATCAGTTGCTCAAGGCATA
TATGATGTCAATATCTCCGGTCTGGTAAGCACAACCATGCAGAATGAAGCCCGTCGTCTGCGTGCCGAACGCTGG
AAAGCGGAAAATCAGGAAGGGATGGCTGAGGTCGCCCGGTTTATTGAAATGAACGGCTCTTTTGCTGACGAGAAC
AGGGGCTGGTGAAATGCAGTTTAAGGTTTACACCTATAAAAGAGAGAGCCGTTATCGTCTGTTTGTGGATGTACA
GAGTGATATTATTGACACGCCCGGCCGACGGATGGTGATCCCCCTGGCCAGTGCACGTCTGCTGTCAGATAAAGT
CTCCCGTGAACTTTACCCGGTGGTGCATATCGGGGATGAAAGCTGGCGCATGATGACCACCGATATGGCCAGTGT
GCCGGTTTCCGTTATCGGGGAAGAAGTGGCTGATCTCAGCCACCGCGAAAATGACATCAAAAACGCCATTAACCT
GATGTTCTGGGGAATATAAATGTCAGGCTCCCTTATACACAGCCAGTCTGCACCTCGACggtctcAcattggctc
ttcttactAAGGGTGGGCGCGCCGACCCAGCTTTCTTGTACAAAGTTGGCATTATAAGAAAGCATTGCTTATCAA
TTTGTTGCAACGAACAGGTCACTATCAGTCAAAATAAAATCATTATTTGCCATCCAGCTGATATCCCCTATAGTG
AGTCGTATTACATGGTCATAGCTGTTTCCTGGCAGCTCTGGCCCGTGTCTCAAAATCTCTGATGTTACATTGCAC
AAGATAAAAATATATCATCATGAACAATAAAACTGTCTGCTTACATAAACAGTAATACAAGGGGTGTTatgagcc
atattcaacgggaaacgtcgaggccgcgattaaattccaacatggatgctgatttatatgggtataaatgggctc
gcgataatgtcgggcaatcaggtgcgacaatctatcgcttgtatgggaagcccgatgcgccagagttgtttctga
aacatggcaaaggtagcgttgccaatgatgttacagatgagatggtcagactaaactggctgacggaatttatgc
ctcttccgaccatcaagcattttatccgtactcctgatgatgcatggttactcaccactgcgatccccggaaaaa
cagcattccaggtattagaagaatatcctgattcaggtgaaaatattgttgatgcgctggcagtgttcctgcgcc
ggttgcattcgattcctgtttgtaattgtccttttaacagcgatcgcgtatttcgtctcgctcaggcgcaatcac
gaatgaataacggtttggttgatgcgagtgattttgatgacgagcgtaatggctggcctgttgaacaagtctgga
aagaaatgcataaacttttgccattctcaccggattcagtcgtcactcatggtgatttctcacttgataacctta
tttttgacgaggggaaattaataggttgtattgatgttggacgagtcggaatcgcagaccgataccaggatcttg
ccatcctatggaactgcctcggtgagttttctccttcattacagaaacggctttttcaaaaatatggtattgata
atcctgatatgaataaattgcagtttcatttgatgctcgatgagtttttcTAATCAGAATTGGTTAATTGGTTGT
AACACTGGCAGAGCATTACGCTGACTTGACGGGACGGCGCAAGCTCATGACCAAAATCCCTTAACGTGAGTTACG
CGTCGTTCCACTGAGCGTCAGACCCCGTAGAAAAGATCAAAGGATCTTCTTGAGATCCTTTTTTTCTGCGCGTAA
TCTGCTGCTTGCAAACAAAAAAACCACCGCTACCAGCGGTGGTTTGTTTGCCGGATCAAGAGCTACCAACTCTTT
TTCCGAAGGTAACTGGCTTCAGCAGAGCGCAGATACCAAATACTGTCCTTCTAGTGTAGCCGTAGTTAGGCCACC
ACTTCAAGAACTCTGTAGCACCGCCTACATACCTCGCTCTGCTAATCCTGTTACCAGTGGCTGCTGCCAGTGGCG
ATAAGTCGTGTCTTACCGGGTTGGACTCAAGACGATAGTTACCGGATAAGGCGCAGCGGTCGGGCTGAACGGGGG
GTTCGTGCACACAGCCCAGCTTGGAGCGAACGACCTACACCGAACTGAGATACCTACAGCGTGAGCATTGAGAAA
GCGCCACGCTTCCCGAAGGGAGAAAGGCGGACAGGTATCCGGTAAGCGGCAGGGTCGGAACAGGAGAGCGCACGA
GGGAGCTTCCAGGGGGAAACGCCTGGTATCTTTATAGTCCTGTCGGGTTTCGCCACCTCTGACTTGAGCGTCGAT

TTTTGTGATGCTCGTCAGGGGGGCGGAGCCTATGGAAAAACGCCAGCAACGCGGCCTTTTTACGGTTCCTGGCCT
TTTGCTGGCCTTTTGCTCACATGTT

PURPLE/UPPERCASE: M13-F binding site
orange/lowercase: attL1
BLUE/UPPERCASE: *AtMIR390a* 5' region
**RED/UPPERCASE/BOLD:** A18G mutation
RED/UPPERCASE: *Bsa*I site
magenta/lowercase: chloramphenicol resistance gene
MAGENTA/UPPERCASE: *ccd*B gene
red/lowercase: inverted *Bsa*I site
blue/lowercase: *AtMIR390a* 3' region
orange/lowercase/underlined: attL2
PURPLE/UPPERCASE/UNDERLINED: M13-Reverse binding site
brown/lowercase: Kanamycin resistance gene

**>*pMDC32B-BS-AtMIR390-A18G-B/c* (11629 bp)**

CCAGCCAGCCAACAGCTCCCCGACCGGCAGCTCGGCACAAAATCACCACTCGATACAGGCAGCCCATCAGTCCGG
GACGGCGTCAGCGGGAGAGCCGTTGTAAGGCGGCAGACTTTGCTCATGTTACCGATGCTATTCGGAAGAACGGCA
ACTAAGCTGCCGGGTTTGAAACACGGATGATCTCGCGGAGGGTAGCATGTTGATTGTAACGATGACAGAGCGTTG
CTGCCTGTGATCACCGCGGTTTCAAAATCGGCTCCGTCGATACTATGTTATACGCCAACTTTGAAAACAACTTTG
AAAAAGCTGTTTTCTGGTATTTAAGGTTTTAGAATGCAAGGAACAGTGAATTGGAGTTCGTCTTGTTATAATTAG
CTTCTTGGGGTATCTTTAAATACTGTAGAAAAGAGGAAGGAAATAATAAatggctaaaatgagaatatcaccgga
attgaaaaaactgatcgaaaaataccgctgcgtaaaagatacggaaggaatgtctcctgctaaggtatataagct
ggtgggagaaaatgaaaacctatatttaaaaatgacggacagccggtataaagggaccacctatgatgtggaacg
ggaaaaggacatgatgctatggctggaaggaaagctgcctgttccaaaggtcctgcactttgaacggcatgatgg
ctggagcaatctgctcatgagtgaggccgatggcgtcctttgctcggaagagtatgaagatgaacaaagccctga
aaagattatcgagctgtatgcggagtgcatcaggctctttcactccatcgacatatcggattgtccctatacgaa
tagcttagacagccgcttagccgaattggattacttactgaataacgatctggccgatgtggattgcgaaaactg
ggaagaagacactccatttaaagatccgcgcgagctgtatgatttttttaaagacggaaaagcccgaagaggaact
tgtcttttcccacggcgacctgggagacagcaacatctttgtgaaagatggcaaagtaagtggctttattgatct
tgggagaagcggcagggcggacaagtggtatgacattgccttctgcgtccggtcgatcagggaggatatcgggga
agaacagtatgtcgagctatttttttgacttactggggatcaagcctgattgggagaaaataaaatattatattt
actggatgaattgttttagTACCTAGAATGCATGACCAAAATCCCTTAACGTGAGTTTTCGTTCCACTGAGCGTC
AGACCCCGTAGAAAAGATCAAAGGATCTTCTTGAGATCCTTTTTTTCTGCGCGTAATCTGCTGCTTGCAAACAAA
AAAACCACCGCTACCAGCGGTGGTTTGTTTGCCGGATCAAGAGCTACCAACTCTTTTTCCGAAGGTAACTGGCTT
CAGCAGAGCGCAGATACCAAATACTGTCCTTCTAGTGTAGCCGTAGTTAGGCCACCACTTCAAGAACTCTGTAGC
ACCGCCTACATACCTCGCTCTGCTAATCCTGTTACCAGTGGCTGCTGCCAGTGGCGATAAGTCGTGTCTTACCGG
GTTGGACTCAAGACGATAGTTACCGGATAAGGCGCAGCGGTCGGGCTGAACGGGGGGTTCGTGCACACAGCCCAG
CTTGGAGCGAACGACCTACACCGAACTGAGATACCTACAGCGTGAGCTATGAGAAAGCGCCACGCTTCCCGAAGG
GAGAAAGGCGGACAGGTATCCGGTAAGCGGCAGGGTCGGAACAGGAGAGCGCACGAGGGAGCTTCCAGGGGGAAA
CGCCTGGTATCTTTATAGTCCTGTCGGGTTTCGCCACCTCTGACTTGAGCGTCGATTTTTGTGATGCTCGTCAGG
GGGGCGGAGCCTATGGAAAAACGCCAGCAACGCGGCCTTTTTACGGTTCCTGGCCTTTTGCTGGCCTTTTGCTCA
CATGTTCTTTCCTGCGTTATCCCCTGATTCTGTGGATAACCGTATTACCGCCTTTGAGTGAGCTGATACCGCTCG
CCGCAGCCGAACGACCGAGCGCAGCGAGTCAGTGAGCGAGGAAGCGGAAGAGCGCCTGATGCGGTATTTTCTCCT
TACGCATCTGTGCGGTATTTCACACCGCATATGGTGCACTCTCAGTACAATCTGCTCTGATGCCGCATAGTTAAG
CCAGTATACACTCCGCTATCGCTACGTGACTGGGTCATGGCTGCGCCCCGACACCCGCCAACACCCGCTGACGCG
CCCTGACGGGCTTGTCTGCTCCCGGCATCCGCTTACAGACAAGCTGTGACCGTCTCCGGGAGCTGCATGTGTCAG
AGGTTTTCACCGTCATCACCGAAACGCGCGAGGCAGGGTGCCTTGATGTGGGCGCCGGCGGTCGAGTGGCGACGG
CGCGGCTTGTCCGCGCCCTGGTAGATTGCCTGGCCGTAGGCCAGCCATTTTTGAGCGGCCAGCGGCCGCGATAGG
CCGACGCGAAGCGGCGGGGCGTAGGGAGCGCAGCGACCGAAGGGTAGGCGCTTTTTGCAGCTCTTCGGCTGTGCG
CTGGCCAGACAGTTATGCACAGGCCAGGCGGGTTTTAAGAGTTTTAATAAGTTTTAAAGAGTTTTAGGCGGAAAA
ATCGCCTTTTTTCTCTTTTATATCAGTCACTTACATGTGTGACCGGTTCCCAATGTACGGCTTTGGGTTCCCAAT
GTACGGGTTCCGGTTCCCAATGTACGGCTTTGGGTTCCCAATGTACGTGCTATCCACAGGAAAGAGAACTTTTCG
ACCTTTTTCCCCTGCTAGGGCAATTTGCCCTAGCATCTGCTCCGTACATTAGGAACCGGCGGATGCTTCGCCCTC
GATCAGGTTGCGGTAGCGCATGACTAGGATCGGGCCAGCCTGCCCCGCCTCCTCCTTCAAATCGTACTCCGGCAG
GTCATTTGACCCGATCAGCTTGCGCACGGTGAAACAGAACTTCTTGAACTCTCCGGCGCTGCCACTGCGTTCGTA
GATCGTCTTGAACAACCATCTGGCTTCTGCCTTGCCTGCGGCGCGGCGTGCCAGGCGGTAGAGAAAACGGCCGAT
GCCGGGATCGATCAAAAAGTAATCGGGGTGAACCGTCAGCACGTCCGGGTTCTTGCCTTCTGTGATCTCGCGGTA
CATCCAATCAGCTAGCTCGATCTCGATGTACTCCGGCCGCCCGGTTTCGCTCTTTACGATCTTGTAGCGGCTAAT
CAAGGCTTCACCCTCGGATACCGTCACCAGGCGGCCGTTCTTGGCCTTCTTCGTACGCTGCATGGCAACGTGCGT
GGTGTTTAACCGAATGCAGGTTTCTACCAGGTCGTCTTTCTGCTTTCCGCCATCGGCTCGCCGGCAGAACTTGAG
TACGTCCGCAACGTGTGGACGGAACACGCGGCCGGGCTTGTCTCCCTTCCCTTCCCGGTATCGGTTCATGGATTC
GGTTAGATGGGAAACCGCCATCAGTACCAGGTCGTAATCCCACACACTGGCCATGCCGGCCGGCCCTGCGGAAAC
CTCTACGTGCCCGTCTGGAAGCTCGTAGCGGATCACCTCGCCAGCTCGTCGGTCACGCTTCGACAGACGGAAAAC
GGCCACGTCCATGATGCTGCGACTATCGCGGGTGCCCACGTCATAGAGCATCGGAACGAAAAAATCTGGTTGCTC
GTCGCCCTTGGGCGGCTTCCTAATCGACGGCGCACCGGCTGCCGGCGGTTGCCGGGATTCTTTGCGGATTCGATC
AGCGGCCGCTTGCCACGATTCACCGGGGCGTGCTTCTGCCTCGATGCGTTGCCGCTGGGCGGCCTGCGCGGCCTT
CAACTTCTCCACCAGGTCATCACCCAGCGCCGCGCCGATTTGTACCGGGCCGGATGGTTTGCGACCGTCACGCCG
ATTCCTCGGGCTTGGGGGTTCCAGTGCCATTGCAGGGCCGGCAGACAACCCAGCCGCTTACGCCTGGCCAACCGC
CCGTTCCTCCACACATGGGGCATTCCACGGCGTCGGTGCCTGGTTGTTCTTGATTTTCCATGCCGCCTCCTTTAG
CCGCTAAAATTCATCTACTCATTTATTCATTTGCTCATTTACTCTGGTAGCTGCGCGATGTATTCAGATAGCAGC
TCGGTAATGGTCTTGCCTTGGCGTACCGCGTACATCTTCAGCTTGGTGTGATCCTCCGCCGGCAACTGAAAGTTG
ACCCGCTTCATGGCTGGCGTGTCTGCCAGGCTGGCCAACGTTGCAGCCTTGCTGCTGCGTGCGCTCGGACGGCCG
GCACTTAGCGTGTTTGTGCTTTTGCTCATTTTCTCTTTACCTCATTAACTCAAATGAGTTTTGATTTAATTTCAG
CGGCCAGCGCCTGGACCTCGCGGGCAGCGTCGCCCTCGGGTTCTGATTCAAGAACGGTTGTGCCGGCGGCGGCAG
TGCCTGGGTAGCTCACGCGCTGCGTGATACGGGACTCAAGAATGGGCAGCTCGTACCCGGCCAGCGCCTCGGCAA

CCTCACCGCCGATGCGCGTGCCTTTGATCGCCCGCGACACGACAAAGGCCGCTTGTAGCCTTCCATCCGTGACCT
CAATGCGCTGCTTAACCAGCTCCACCAGGTCGGCGGTGGCCCATATGTCGTAAGGGCTTGGCTGCACCGGAATCA
GCACGAAGTCGGCTGCCTTGATCGCGGACACAGCCAAGTCCGCCGCCTGGGGCGCTCCGTCGATCACTACGAAGT
CGCGCCGGCCGATGGCCTTCACGTCGCGGTCAATCGTCGGGCGGTCGATGCCGACAACGGTTAGCGGTTGATCTT
CCCGCACGGCCGCCCAATCGCGGGCACTGCCCTGGGGATCGGAATCGACTAACAGAACATCGGCCCCGGCGAGTT
GCAGGGCGCGGGCTAGATGGGTTGCGATGGTCGTCTTGCCTGACCCGCCTTTCTGGTTAAGTACAGCGATAACCT
TCATGCGTTCCCCTTGCGTATTTGTTTATTTACTCATCGCATCATATACGCAGCGACCGCATGACGCAAGCTGTT
TTACTCAAATACACATCACCTTTTTAGACGGCGGCGCTCGGTTTCTTCAGCGGCCAAGCTGGCCGGCCAGGCCGC
CAGCTTGGCATCAGACAAACCGGCCAGGATTTCATGCAGCCGCACGGTTGAGACGTGCGCGGGCGGCTCGAACAC
GTACCCGGCCGCGATCATCTCCGCCTCGATCTCTTCGGTAATGAAAAACGGTTCGTCCTGGCCGTCCTGGTGCGG
TTTCATGCTTGTTCCTCTTGGCGTTCATTCTCGGCGGCCGCCAGGGCGTCGGCCTCGGTCAATGCGTCCTCACGG
AAGGCACCGCGCCGCCTGGCCTCGGTGGGCGTCACTTCCTCGCTGCGCTCAAGTGCGCGGTACAGGGTCGAGCGA
TGCACGCCAAGCAGTGCAGCCGCCTCTTTCACGGTGCGGCCTTCCTGGTCGATCAGCTCGCGGGCGTGCGCGATC
TGTGCCGGGGTGAGGGTAGGGCGGGGGCCAAACTTCACGCCTCGGGCCTTGGCGGCCTCGCGCCCGCTCCGGGTG
CGGTCGATGATTAGGGAACGCTCGAACTCGGCAATGCCGGCGAACACGGTCAACACCATGCGGCCGGCCGGCGTG
GTGGTGTCGGCCCACGGCTCTGCCAGGCTACGCAGGCCCGCGCCGGCCTCCTGGATGCGCTCGGCAATGTCCAGT
AGGTCGCGGGTGCTGCGGGCCAGGCGGTCTAGCCTGGTCACTGTCACAACGTCGCCAGGGCGTAGGTGGTCAAGC
ATCCTGGCCAGCTCCGGGCGGTCGCGCCTGGTGCCGGTGATCTTCTCGGAAAACAGCTTGGTGCAGCCGGCCGCG
TGCAGTTCGGCCCGTTGGTTGGTCAAGTCCTGGTCGTCGGTGCTGACGCGGGCATAGCCCAGCAGGCCAGCGGCG
GCGCTCTTGTTCATGGCGTAATGTCTCCGGTTCTAGTCGCAAGTATTCTACTTTATGCGACTAAAACACGCGACA
AGAAAACGCCAGGAAAAGGGCAGGGCGGCAGCCTGTCGCGTAACTTAGGACTTGTGCGACATGTCGTTTTCAGAA
GACGGCTGCACTGAACGTCAGAAGCCGACTGCACTATAGCAGCGGAGGGGTTGGATCAAAGTACTTTGATCCCGA
GGGGAACCCTGTGGTTGGCATGCACATACAAATGGACGAACGGATAAACCTTTTCACGCCCTTTTAAATATCCGT
TATTCTAATAAACGCTCTTTTCTCTTAGGtttacccgccaatatatcctgtcaAACACTGATAGTTTAAACTGAA
GGCGGGAAACGACAATCTGATCCAAGCTCAAGCTGCTCTAGCATTCGCCATTCAGGCTGCGCAACTGTTGGGAAG
GGCGATCGGTGCGGGCCTCTTCGCTATTACGCCAGCTGGCGAAAGGGGGATGTGCTGCAAGGCGATTAAGTTGGG
TAACGCCAGGGTTTTCCCAGTCACGACGTTGTAAAACGACGGCCAGTGCCAAGCTTGGCGTGCCTGCAGGTCAAC
ATGGTGGAGCACGACACACTTGTCTACTCCAAAAATATCAAAGATACAGTCTCAGAAGACCAAAGGGCAATTGAG
ACTTTTCAACAAAGGGTAATATCCGGAAACCTCCTCGGATTCCATTGCCCAGCTATCTGTCACTTTATTGTGAAG
ATAGTGGAAAAGGAAGGTGGCTCCTACAAATGCCATCATTGCGATAAAGGAAAGGCCATCGTTGAAGATGCCTCT
GCCGACAGTGGTCCCAAAGATGGACCCCCACCCACGAGGAGCATCGTGGAAAAAGAAGACGTTCCAACCACGTCT
TCAAAGCAAGTGGATTGATGTGATAACATGGTGGAGCACGACACACTTGTCTACTCCAAAAATATCAAAGATACA
GTCTCAGAAGACCAAAGGGCAATTGAGACTTTTCAACAAAGGGTAATATCCGGAAACCTCCTCGGATTCCATTGC
CCAGCTATCTGTCACTTTATTGTGAAGATAGTGGAAAAGGAAGGTGGCTCCTACAAATGCCATCATTGCGATAAA
GGAAAGGCCATCGTTGAAGATGCCTCTGCCGACAGTGGTCCCAAAGATGGACCCCCACCCACGAGGAGCATCGTG
GAAAAAGAAGACGTTCCAACCACGTCTTCAAAGCAAGTGGATTGATGTGATATCTCCACTGACGTAAGGGATGAC
GCACAATCCCACTATCCTTCGCAAGACCCTTCCTCTATATAAGGAAGTTCATTTCATTTGGAGAGGACCTCGACT
CTAGAGGATCCCCGGGTACCGGGCCCCCCCTCGAGGCGCGCCAAGCTATCAAACAAGTTTGTACAAAAAAGCAGG
CTCCGCGGCCGCCCCCTTCACCAGTAGAGAAGAATCTGT**G**AGAGACCATTAGGCACCCCAGGCTTTACACTTTAT
GCTTCCGGCTCGTATAATGTGTGGATTTTGAGTTAGGAGCCGTCGAGATTTTCAGGAGCTAAGGAAGCTAAAatg
gagaaaaaaatcactggatataccaccgttgatatatcccaatggcatcgtaaagaacattttgaggcatttcag
tcagttgctcaatgtacctataaccagaccgttcagctggatattacggcctttttaaagaccgtaaagaaaaat
aagcacaagttttatccggcctttattcacattcttgcccgcctgatgaatgctcatccggagttccgtatggca
atgaaagacggtgagctggtgatatgggatagtgttcacccttgttacaccgttttccatgagcaaactgaaacg
ttttcatcgctctggagtgaataccacgacgatttccggcagtttctacacatatattcgcaagatgtggcgtgt
tacggtgaaaacctggcctatttccctaaagggtttattgagaatatgtttttcgtctcagccaatccctgggtg
agtttcaccagttttgatttaaacgtggccaatatggacaacttcttcgcccccgttttcaccatgggcaaatat
tatacgcaaggcgacaaggtgctgatgccgctggcgattcaggttcatcatgccgtttgtgatggcttccatgtc
ggcagaatgcttaatgaattacaacagtactgcgatgagtggcagggcggggcgtaaACGCGTGGAGCCGGCTTA
CTAAAAGCCAGATAACAGTATGCGTATTTGCGCGCTGATTTTTGCGGTATAAGAATATATACTGATATGTATACC
CGAAGTATGTCAAAAAGAGGTATGCTATGAAGCAGCGTATTACAGTGACAGTTGACAGCGACAGCTATCAGTTGC
TCAAGGCATATATGATGTCAATATCTCCGGTCTGGTAAGCACAACCATGCAGAATGAAGCCCGTCGTCTGCGTGC
CGAACGCTGGAAAGCGGAAAATCAGGAAGGGATGGCTGAGGTCGCCCGGTTTATTGAAATGAACGGCTCTTTTGC
TGACGAGAACAGGGGCTGGTGAAATGCAGTTTAAGGTTTACACCTATAAAAGAGAGAGCCGTTATCGTCTGTTTG
TGGATGTACAGAGTGATATTATTGACACGCCCGGCCGACGGATGGTGATCCCCCTGGCCAGTGCACGTCTGCTGT
CAGATAAAGTCTCCCGTGAACTTTACCCGGTGGTGCATATCGGGGATGAAAGCTGGCGCATGATGACCACCGATA
TGGCCAGTGTGCCGGTTTCCGTTATCGGGGAAGAAGTGGCTGATCTCAGCCACCGCGAAAATGACATCAAAAACG
CCATTAACCTGATGTTCTGGGGAATATAAATGTCAGGCTCCCTTATACACAGCCAGTCTGCACCTCGACggtctc
AcattggctcttcttactAAGGGTGGGCGCGCCGACCCAGCTTTCTTGTACAAAGTGGTTCGATAATTCCTTAAT
TAACTAGTTCTAGAGCGGCCGCCCACCGCGGTGGAGCTCGAATTTCCCCGATCGTTCAAACATTTGGCAATAAAG
TTTCTTAAGATTGAATCCTGTTGCCGGTCTTGCGATGATTATCATATAATTTCTGTTGAATTACGTTAAGCATGT

AATAATTAACATGTAATGCATGACGTTATTTATGAGATGGGTTTTTATGATTAGAGTCCCGCAATTATACATTTA
ATACGCGATAGAAAACAAAATATAGCGCGCAAACTAGGATAAATTATCGCGCGCGGTGTCATCTATGTTACTGAA
TTCGTAATCATGGTCATAGCTGTTTCCTGTGTGAAATTGTTATCCGCTCACAATTCCACACAACATACGAGCCGG
AAGCATAAAGTGTAAAGCCTGGGGTGCCTAATGAGTGAGCTAACTCACATTAATTGCGTTGCGCTCACTGCCCGC
TTTCCAGTCGGGAAACCTGTCGTGCCAGCTGCATTAATGAATCGGCCAACGCGCGGGGAGAGGCGGTTTGCGTAT
TGGCTAGAGCAGCTTGCCAACATGGTGGAGCACGACACTCTCGTCTACTCCAAGAATATCAAAGATACAGTCTCA
GAAGACCAAAGGGCTATTGAGACTTTTCAACAAAGGGTAATATCGGGAAACCTCCTCGGATTCCATTGCCCAGCT
ATCTGTCACTTCATCAAAAGGACAGTAGAAAAGGAAGGTGGCACCTACAAATGCCATCATTGCGATAAAGGAAAG
GCTATCGTTCAAGATGCCTCTGCCGACAGTGGTCCCAAAGATGGACCCCCACCCACGAGGAGCATCGTGGAAAAA
GAAGACGTTCCAACCACGTCTTCAAAGCAAGTGGATTGATGTGATAACatggtggagcacgacactctcgtctac
tccaagaatatcaaagatacagtctcagaagaccaaagggctattgagacttttcaacaaagggtaatatcggga
aacctcctcggattccattgcccagctatctgtcacttcatcaaaaggacagtagaaaaggaaggtggcacctac
aaatgccatcattgcgataaaggaaaggctatcgttcaagatgcctctgccgacagtggtcccaaagatggaccc
ccacccacgaggagcatcgtggaaaaagaagacgttccaaccacgtcttcaaagcaagtggattgatgtgatatc
tccactgacgtaagggatgacgcacaatcccactatccttcgcaagaccttcctctatataaggaagttcatttc
atttggagaggACACGCTGAAATCACCAGTCTCTCTCTACAAATCTATCTCTCTCGAGCTTTCGCAGATCCCGGG
GGGCAATGAGATATGAAAAAGCCTGAACTCACCGCGACGTCTGTCGAGAAGTTTCTGATCGAAAAGTTCGACAGC
GTCTCCGACCTGATGCAGCTCTCGGAGGGCGAAGAATCTCGTGCTTTCAGCTTCGATGTAGGAGGGCGTGGATAT
GTCCTGCGGGTAAATAGCTGCGCCGATGGTTTCTACAAAGATCGTTATGTTTATCGGCACTTTGCATCGGCCGCG
CTCCCGATTCCGGAAGTGCTTGACATTGGGGAGTTTAGCGAGAGCCTGACCTATTGCATCTCCCGCCGTGCACAG
GGTGTCACGTTGCAAGACCTGCCTGAAACCGAACTGCCCGCTGTTCTACAACCGGTCGCGGAGGCTATGGATGCG
ATCGCTGCGGCCGATCTTAGCCAGACGAGCGGGTTCGGCCCATTCGGACCGCAAGGAATCGGTCAATACACTACA
TGGCGTGATTTCATATGCGCGATTGCTGATCCCCATGTGTATCACTGGCAAACTGTGATGGACGACACCGTCAGT
GCGTCCGTCGCGCAGGCTCTCGATGAGCTGATGCTTTGGGCCGAGGACTGCCCCGAAGTCCGGCACCTCGTGCAC
GCGGATTTCGGCTCCAACAATGTCCTGACGGACAATGGCCGCATAACAGCGGTCATTGACTGGAGCGAGGCGATG
TTCGGGGATTCCCAATACGAGGTCGCCAACATCTTCTTCTGGAGGCCGTGGTTGGCTTGTATGGAGCAGCAGACG
CGCTACTTCGAGCGGAGGCATCCGGAGCTTGCAGGATCGCCACGACTCCGGGCGTATATGCTCCGCATTGGTCTT
GACCAACTCTATCAGAGCTTGGTTGACGGCAATTTCGATGATGCAGCTTGGGCGCAGGGTCGATGCGACGCAATC
GTCCGATCCGGAGCCGGGACTGTCGGGCGTACACAAATCGCCCGCAGAAGCGCGGCCGTCTGGACCGATGGCTGT
GTAGAAGTACTCGCCGATAGTGGAAACCGACGCCCCAGCACTCGTCCGAGGGCAAAGAATAGAGTAGATGCCGA
CCGGATCTGTCGATCGACAAGCTCGAGtttctccataataatgtgtgagtagttcccagataagggaattagggt
tcctatagggtttcgctcatgtgttgagcatataagaaacccttagtatgtatttgtatttgtaaaatacttcta
tcaataaaatttctaattcctaaaaccaaaatccagtactaaaatccagatcCCCCGAATTAATTCGGCGTTAAT
TCAGTACATTAAAAACGTCCGCAATGTGTTATTAAGTTGTCTAAGCGTCAATTTGTTTACACCACAATATATCCT
GCCA

brown/lowercase: kanamycin resistance gene
CYAN/UPPERCASE/UNDERLINED: C->A transversion to block vector's *Bsa*I site
cyan/lowercase: T-DNA right border
GREEN/UPPERCASE: 2x35S CaMV promoter
ORANGE/UPPERCASE: attB1
BLUE/UPPERCASE: *AtMIR390a* 5' region
**RED/UPPERCASE/BOLD:** A18G mutation
RED/UPPERCASE: *Bsa*I site
magenta/lowercase: chloramphenicol resistance gene
MAGENTA/UPPERCASE: *ccd*B gene
red/lowercase: inverted *Bsa*I site
blue/lowercase: *AtMIR390a* 3' region
ORANGE/UPPERCASE/UNDERLINED: attB2
GREY/UPPERCASE/UNDERLINED: Nos terminator
green/lowercase: CaMV promoter
BROWN/UPPERCASE: hygromycin resistance gene
green/lowercase/underlined: CaMV terminator
CYAN/UPPERCASE: T-DNA left border
